# Molecular Architecture and Function Mechanism of Tri-heteromeric GluN1-N2-N3A NMDA Receptors

**DOI:** 10.1101/2025.08.25.672209

**Authors:** Zengwei Kou, Fenyong Yao, Tongtong Zhang, Nan Song, Chun Xie, Boshuang Wang, Yidi Sun

**Author notes:** **Correspondence**: Dr. Zengwei Kou. These authors contributed equally (Zengwei Kou, Fenyong Yao).

## Abstract

*N*-methyl-D-aspartate receptors (NMDARs) play a pivotal role in brain development and synaptic function. Previous studies have focused on GluN1-N2 (2A-2D) and GluN1- N3 (3A and 3B) di-heteromeric (di-) NMDARs, leaving the activation mechanism and stoichiometry of GluN1-N2-N3 tri-heteromeric (tri-) NMDARs largely unexplored. In this study, we employed a two-step affinity-tagged chromatography approach to purify recombinantly expressed GluN1-N2A-N3A tri-NMDARs and determined their cryo-EM structure. Based on the proteoliposome single-channel recording, we discovered GluN1-N2A-N3A can be activated upon co-binding of glycine and glutamate, exhibiting two distinct conductance levels. Furthermore, leveraging structural-based click-chemistry, we introduced photo-crosslinker *p*-azido-phenylalanine (AzF) into the N-terminal domain of GluN2A and GluN2B, enabling the crosslinking with GluN3A subunit both *in vitro* and *in vivo*. These findings provide molecular insights into the subunit arrangement, native architecture and activation mechanism of GluN1-N2-N3A tri- NMDARs and also highlight the complexity of NMDAR assembly and function in the brain.

## Main

NMDARs function as coincidence detectors in neural signaling, typically requiring both membrane depolarization (to relieve Mg^2+^ block) and the binding of co-agonists glycine and glutamate for the opening of ion channel gate. Upon activation, they permit the influx of Na^+^ and Ca^2+^, playing a critical role in synaptic plasticity, learning and memory. In mammals, NMDARs are encoded by seven genes, comprising glycine- bound GluN1 and GluN3 (3A and 3B) subunits, and glutamate-bound GluN2 (from N2A to N2D) subunits. Synaptic NMDARs typically assemble as di- or tri-heterotetramers, consisting of two obligatory GluN1 subunits and two alternative subunits selectively chosen from GluN2 (2A-2D) ^1–8^ and/or GluN3 (3A-3B) subunits^9–11^. Different receptor subtypes exhibit distinct biophysical and pharmacological properties, which are primarily determined by subtle structural variations in their individual domains^7,12^. For instance, the activation of GluN1-N2 di-NMDARs requires the presence of both glycine and glutamate agonists, while GluN1-N3 di-NMDARs only necessitate glycine. Furthermore, the GluN1-N2 subtypes demonstrate greater Ca^2+^ permeability, and enhanced Mg^2+^ sensitivity in contrast to GluN1-N3 subtypes. Additionally, classic GluN1-N2 di-NMDARs pore blocker like ketamine, memantine, showed only small effect to GluN1-N3 di-NMDARs.

Differing from the conventional GluN1-N2 receptors, GluN3A-containing NMDARs exhibit unique characteristics in their expression profiles, structural arrangement^13–15^ physiological and pathological functions. In mice, the GluN3A subunit displays a distinct temporal and spatial expression pattern, showing widespread brain expression during early development that peaks between postnatal days 7-14^16,17^, followed by a decline after adolescence ^18–20^. However, it maintain relatively high expression levels in specific adult brain regions, including the neocortex S1 region^21^, visual cortex layer^22^, basolateral amygdala (BLA)^23^,and media habenula (MHb)^24^. During brain development, GluN3A is suggested serve a dominant negative role by suppressing neuronal over- development tendency through decreasing synaptic stability and promoting the pruning of superfluous spines^19,25^. Additionally, it regulates multiple synaptic events including the synaptic trafficking of GluN2A-containing NMDARs^26^, pre-synaptic release of neurotransmitter^27^ and NMDAR-induced transcription^28^. Through modulation of astrocyte-neuron crosstalk, neocortical circuits, and synaptic plasticity, GluN3A- containing NMDARs influence visual^22^, and learning processes^19^, as well as emotional and social behavior^23,24^. Notably, abnormalities in GluN3A expression and function have been associated with various neurological brain disorders including addiction^29,30^, ischemia brain stroke^31–34^, Huntington’s disease^35–37^, Alzheimer’s disease^38^, addiction^29,39^, epilepsy^40^, autoimmune encephalitis^41^, and depression^42^.

Prior studies using recombinant expression systems have demonstrated that GluN1-N3A di-NMDARs exhibit lower Ca^2+^ permeability, less sensitive to Mg^2+^ blockage, and faster desensitization kinetics^43–45^, compared to the conventional GluN1-N2 di- NMDARs. Moreover, the activation mechanisms of GluN1-N3A di-NMDARs differ fundamentally from that of GluN1-N2 di-NMDARs. Specifically, glycine binding to high-affinity GluN3A sites activates the ion channel, while binding to low-affinity GluN1 sites results in channel to desensitize or pre-desensitize of the channel. When glycine simultaneously binds to both GluN1 and GluN3A subunits, it triggers a rapid conformational change termed as the “apo-open-desensitization” transition^46^. Therefore, blocking glycine binding to GluN1 with competitive antagonists (CGP-78608^47^ / L- 689,560^48^ / MDL-29951^44,49^) or introducing mutations^44,50^ enhances current amplitude and slows desensitization in GluN1-N3A di-NMDARs. This approach enables reliable recordings of native GluN1-N3A di-NMDAR currents in the brain^23,24,51^.

Other studies have also proposed the existence of tri-NMDARs, consisting of two different GluN2 subunits^5,7,52^ or one copy of GluN2 and one copy of GluN3 subunit^11,39^. In recombinant system expressing GluN1 and GluN2A subunits, GluN3A overexpression has been shown to reduce single-channel conductance and Ca^2+^ permeability^53–55^, findings further confirmed in hippocampal neurons^56^. Notably, the presence of GluN1-N2B-N3A tri-NMDARs has been implicated in the expression of cocaine-evoked plasticity ^39^, as well as neocortical synaptic transmission and plasticity^27^. Additionally, GluN3A subunit has been found to co-localized with GluN2A in the postsynaptic regions^11^, and to co-immunoprecipitated with GluN2A^19,26,45,57^, GluN2B^19,45,54,57^, and GluN2C^55^ subunits. However, distinguishing GluN1-N2-N3A tri- NMDARs from GluN1-N2A and GluN1-N3A di-NMDARs remains challenging due to the lack of specific pharmacological and definitive biochemical markers. Consequently, direct evidence for native assembly of GluN3A with GluN2 subunits into GluN1-N2-N3A tri-NMDARs complexes remain elusive. Over the past decades, substantial progress has been made in understanding the molecular basis of agonist recognition and gating mechanisms of GluN1-N2 di- and tri- NMDARs. These receptors assemble as tetrameric complexes with a bouquet-like architecture, comprising the clamshell-shaped extracellular domains of amino-terminal domain (NTD) and ligand-binding domain (LBD), the hydrophobic transmembrane domain (TMD), and the flexible intracellular carboxyl-terminal domain (CTD). Upon binding glutamate and glycine, the LBD clamshell undergoes closure, extending the LBD-TMD linker, which, in turn, imitate gate opening^58^.Notably, GluN1-N3A di-NMDARs showed distinct activate cycle. Glycine binds to GluN3A with high-affinity and this binding activates the ion channel, while glycine binds to GluN1 with low-affinity and this binding leads to desensitization of the channel. The NTD of GluN3A subunit showed highly flexible^13^ and the LBD domain of GluN3A adopts a similar yet denser conformation in the presence of agonist^17,59^. Fundamental questions regarding the subunit stoichiometry, structural arrangement and gating property of GluN1-N2-N3A tri-NMDARs remain unresolved

In this study, we employed a two-step affinity-tagged chromatography purification strategy to determine the cryo-EM structure of GluN1-N2A-N3A tri-NMDARs, revealing an asymmetric tetrameric assembly with a “N1-N2A-N1-N3A” subunit arrangement. Subsequently, we reconstituted the protein into liposomes and conducted single-channel recordings. Our results demonstrated that GluN1-N2A-N3A tri-NMDARs can be activated by the co-application of glycine and glutamate, neither glycine nor glutamate alone. Moreover, the incorporation of the GluN2A subunit appears to play a dominant role in gating, as GluN1-N2A-N3A tri-NMDARs did not exhibit the pronounced glycine-induced desensitization observed in GluN1-N3A di-NMDARs, instead displaying activation properties more akin to GluN1-N2A receptors.

Using genetic code expansion, we introduced a clickable unnatural amino acid AzF (*p*-azido-l-phenylalanine) into the NTD of tri-NMDARs and confirmed GluN2 and GluN3A subunits can be crosslinked in close proximity in the recombinant expression system. Moreover, this click-chemistry strategy enabled us to probe the existence and architecture of these tri-NMDARs in cultured hippocampal neurons and brain tissue. Our findings provide direct evidence that GluN3A can assemble with GluN2 subunits to form functional tri-NMDARs. This work established a framework for structurally and functionally probing multimeric protein complex in native tissues.

## Results

### GluN3A co-expresses with GluN2A/N2B subunits in the brain

We assessed the expression levels of GluN3A and its co-expression patterns with other NMDAR subunits across various brain regions based on previously reported datasets of single-cell RNA sequencing (scRNA-seq) ^60–63^ (Fig. 1a). Our findings revealed significant mRNA expression of GluN3A (*Grin3a*) in excitatory neurons, inhibitory neurons, and non-neuronal cells of the cortex and hippocampus (Extended Data Fig. 1). These results were also supported by the immunohistochemical staining of GluN3A in brain slices and cultured hippocampal neurons (Extended Data Fig.2).

**Figure 1.**
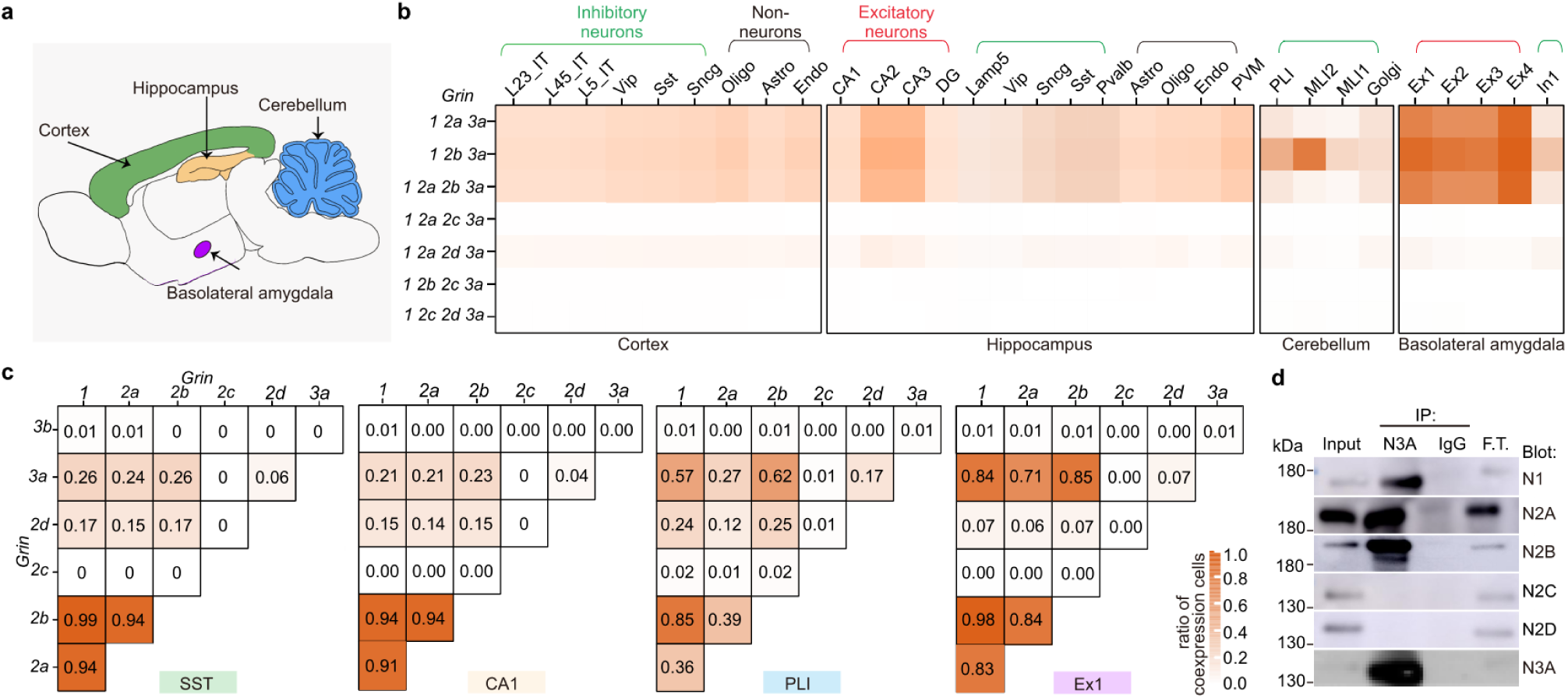
ScRNA-seq and immunoprecipitation analysis of NMDAR subunits co-expression and co-assembly patterns in the brain. **a**. Cartoon illustration depicting the brain regions analyzed in scRNA-seq and Immunoprecipitation. **b**. Co- expression analysis of *Grin3a* with other NMDAR subunits in the cortex, hippocampus, cerebellum, and basolateral amygdala. Colors represent the proportion of cells within each subclass expressing specific subunit combinations. **c**. Paired co-expression analysis of NMDAR subunits in cortex SST neuron, hippocampus CA1 neurons, cerebellum PLI neuron, and basolateral amygdala EX1 neurons, presented as heatmaps. Colors indicate the proportion of cells co-expressing indicated pairs of NMDAR subunits. **d**. Immunoprecipitation (IP) of GluN3A (N3A) with other NMDAR subunits. Total protein from the cortex and hippocampus was precipitated with anti-N3A and subsequently examined by Western blotting using antibodies against NMDAR subunits. Normal IgG (IgG) was used as a negative control. Flow-through (F.T.) fractions were examined to verify effective IP.

We conducted co-expression probability analysis, revealing distinct yet overlapping expression patterns among subunits in GluN3A-highly expressing cell types from cerebral cortex, hippocampus, cerebellum, and basolateral amygdala (BLA). GluN3A exhibited a tendency to co-express with GluN1, GluN2A, and GluN2B subunits instead of co-expressed only with GluN1 subunit. The co-expression probabilities of the GluN3A with GluN2A or GluN2B were more than 24% in two types of inhibitory neurons (24% and 26% in SST neurons of cortex and 27% and 62% in PLI of cerebellum). While in the excitatory hippocampus CA1 neurons, the co-expression probabilities were more than 21% (21% and 23% for GluN2A and GluN2B, respectively). Notably, these co-expression probabilities reached 71% and 85% in excitatory Ex1 neuron of BLA (Fig. 1b-c), where the GluN3A containing NMDARs have been reported to participate in the fear memory.

To determine if GluN3A can associate with GluN2 subunits at the protein level, we conducted a co-immunoprecipitation experiment with native brain tissue. A laboratory-generated monoclonal antibody (clone no. 1G4) specifically recognizing the GluN3A subunit (Extended Data Fig. 3) was used to enrich GluN3A-containing NMDARs from the cortex and hippocampus of mouse brains. Our results showed that GluN3A could be precipitated with GluN1, GluN2A, and GluN2B (Fig. 1d), confirming that potential existence of native GluN1-N2A-N3A and GluN1-N2B-N3A tri-NMDARs, consistent with previous studies^19,45,54,57^. However, under our experimental conditions, we were unable to immunoprecipitated GluN2C and GluN2D with GluN3A subunits (Fig. 1d) likely due to the low abundance of these assemblies.

### Isolation strategy and architecture of GluN1-N2A-N3A tri-NMDARs

Given the above results indicate the presence of GluN1-N2-N3A tri-NMDARs in the brain, we next sought to determine the structure of the GluN1-N2A-N3A receptors. We added a 6 x His tag and a Strep tag to the C-terminal domain (CTD) truncated GluN2A and GluN3A subunits, respectively, and co-expressed them with CTD-truncated GluN1 subunit. Then, we employed a two-step affinity-tagged chromatograph to isolate the GluN1-N2A-N3A tri-NMDARs (Fig. 2a-b) as we previously reported for purification of GluN1-N2A-N2C receptors^7^. The separation of GluN1-N2A and GluN1-N3A di- NMDARs from the GluN1-N2A-N3A tri-NMDARs was confirmed through Coomassie blue staining and western blotting analysis (Fig. 2c).

**Figure 2.**
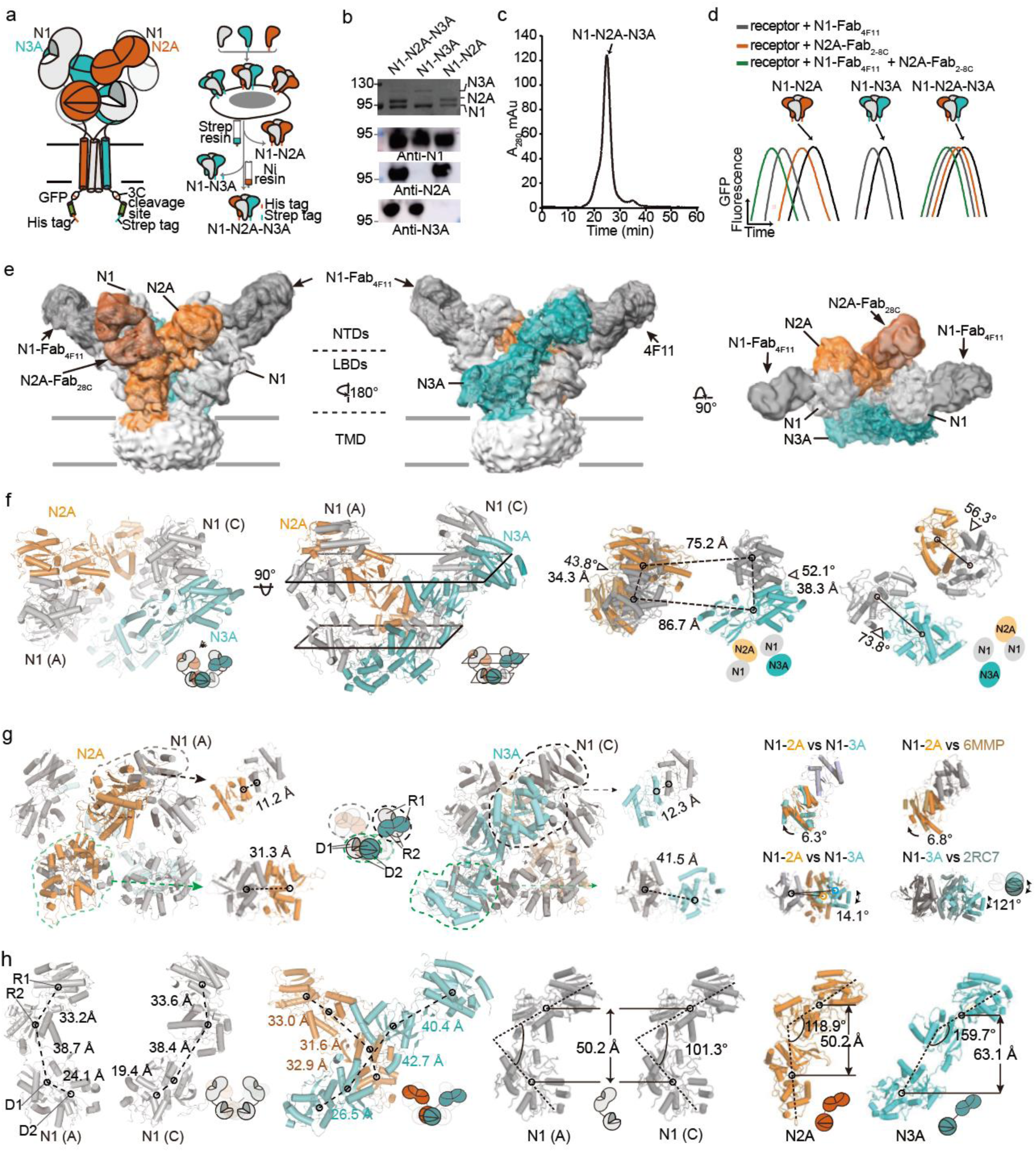
Purification, validation, and structural analysis of N1-N2A-N3A tri- NMDARs. **a**. Topological representation and purification workflow of N1-N2A-N3A. A two-step affinity-tagged chromatographic process was used for the N1-N2A-N3A purification. Fractions containing N1-N2A or N1-N3A are also collected for further analysis. **b**. Coomassie blue staining and Western blot analysis of N1-N2A, N1-N3A, and N1-N2A-N3A receptors from the same preparation batch, post-digestion by Endo H and 3C protease. **c**. SEC traces of N1-N2A- N3A receptors. **d**. FSEC traces depicting the shifts in N1-N2A, N1-N3A, and N1-N2A-N3A receptors upon binding with specific fabs against N1 (clone 4F11), N2A (clone 28C), N3A (clone 1G4), or a combination of fabs against N1 and N2A, as indicated. **e**. Cryo-EM maps of N1-N2A- N3A bound with N1 and N2A Fabs, presented inside and top-down views. The four subunits (N1, N2A, N1, and N3A) are color-coded accordingly as gray, orange, blue, and gray. The N1- specific fab 4F11 is shown in dark gray, and the N2A-specific fab 28C is depicted in brown. **f**. The overall, NTD, and LBD structure of N1-N2A-N3A. **g.** The overall structure of NTD and LBD of N1-N2A and N1-N3A heterodimer. Superimposition of the NTD R1 lobes or LBD from N1- N2A and N1-N3A protomers of N1-2A-3A, or N1-N2A protomers from N1-N2A-N3A and 6MMP. Superimposition of the LBD from N1-N2A protomers of N1-N2A-N3A and crystal N1-N3A LBD structure(2RC7). **h**. Structural analysis of extracellular domains of each protomers. The center- of-mass (COM) of R1 and R2 of NTD, D1 and D2 of LBD of each protomer is indicated by open circles. The distance connected by COMs was measured.

To label the different subunits for cryo-EM classification and reconstruction, we developed two monoclonal antibodies: one bound to the N-terminal domain of GluN1 subunit (clone no. 4F11, referred to as N1-Fab_4F11_)^6^ and another bound to the ligand binding domain (LBD) of GluN2A subunit (clone no. 2-8c, referred to as N2A-Fab_28C_)^5^ (Extended Data Fig.3). We further employed fluorescence-detection size-exclusion chromatography (FSEC) to verify the presence of specific subunits in the assembled NMDAR complexes. GluN1-N2A and GluN1-N2A-N3A receptors were preincubated with N1-Fab_4F11_, N2A-Fab_28C_, or both, while GluN1-N3A receptors were preincubated with N1-Fab_4F11_ alone. Upon Fab binding, the molecular weight of the receptor complex increases, resulting in an earlier elution during size-exclusion chromatography- manifested as a rightward shift in the FSEC profile. For GluN1-N2A-N3A, distinct right shifts were observed upon incubation with both N1-Fab_4F11_ and N2A-Fab_28C_, confirming the incorporation of both GluN1 and GluN2A subunits into the receptor complex (Fig. 2d).

Consequently, we elucidated the structure of GluN1-N2A-N3A tri-NMDARs with N1- Fab_4F11_ and N2A-Fab_28C_ in the presence of agonists glycine and glutamate (Extended Data Fig. 4). In the 2D class averages, we observed the density of N2A-Fab_28C_ protruding from the GluN2A-LBD, along with two N1-Fab_4F11_ densities protruding from the two diagonally opposedGluN1-NTDs (Extended Data Fig. 4b). During 3D classification, we found all four structures contained densities for N2A-Fab_28C_. Two of these structures (accounting for 30.7% of all particles) were better defined and contained two N1-Fab_4F11_ densities. The remaining two less well-defined 3D structures also included a N2A-Fab_28C_ and two N1-Fab_4F11_ at high contour levels. In all structures, only one subunit, the GluN3A subunit, remained Fab free state, with its NTD and LBD domains being less well-defined. Combining the well-defined subclasses, we finally reconstructed the glycine and glutamate-bound form of the GluN1-N2A-N3A tri- NMDARs at an overall resolution of 7.7 Å (Extended Data Fig. 4).

### Subunit arrangement of asymmetric GluN1-N2A-N3A tri-NMDARs

Overall, GluN1-N2A-N3A tri-NMDARs adopt a bouquet-like architecture, similar to conventional GluN1-N2 di-NMDARs^3,4^. The structure resembles an asymmetric dimer-of-dimer organization, where GluN1-N2A and GluN1-N3A heterodimers combine to form a tetrameric assembly. GluN1 subunits occupy positions A and C, while GluN2A and GluN3A subunits occupy positions B and D, respectively. The receptors are organized into three layers: NTD, LBD, and TMD. In the presence of Fabs and agonists, the secondary structural densities for the NTD and LBD remain continuous and defined, enabling rigid-body fitting of R1 and R2 lobes of the NTDs and D1 and D2 lobes of the LBDs into these density features (Fig. 2e, 2f).

In the top-down view, extensive subunit-subunit interactions occur within the NTD, primarily between GluN1-N2A and GluN1-N3A heterodimers, with two GluN1 NTDs positioned at the periphery of the tetrameric assembly. The NTDs of the GluN1 subunit contract with their adjacent GluN2A and GluN3A subunits at the R1 lobes, forming a compressed GluN1-N2A interface and a loose GluN1-N3A interface. The NTD center of mass (COM) distance between GluN1 (protomer A) and the GluN2A subunit is 4.0 Å shorter than that between the GluN1 (protomer C) and the GluN3A subunit, resulting in a COM angle of the GluN1-N2A dimer that is 8.3° smaller than that of the GluN1-N3A dimer. (Fig. 2f).

To gain further insight into the structural asymmetry and subunit-specific interactions within GluN1-N2A-N3A tri-NMDARs, we conduct measurements of the COM distance and vector angle at the NTD and LBD interfaces. At the NTD R1-R1 interface, we observed that the COM distance between α2 of GluN1(A) and α1 of GluN2A is 1.1 Å shorter than that between α2 of GluN1(C) and α1 of GluN3A. Additionally, the vector angle shows 6.3° larger twisting of GluN1 (A) relative to GluN2A compared to GluN1 (C) relative to GluN3A. Notably, this vector angle in the GluN1-N2A -di NMDARs (PDB 6MMP) is 6.8° larger than that in GluN1-N2A-N3A tri-NMDARs.

Importantly, the LBD COM distance between GluN1(C) and GluN2A is 10.2 Å shorter than that between GluN1(A) and GluN3A, with the vector angle exhibiting 14.1° more twisting between GluN1(C)- GluN2A and GluN1(A)-GluN3A.Specifically, the tri-receptor domain shows a 121° anti-clockwise twisting of the whole domain around the pseudo-symmetric axis of the receptor center, which contrasts with the structure of 2RC7 (Fig. 2g, 2h).

We compared two GluN1 subunits in the GluN1-N2A heterodimer and GluN1-N3A heterodimer of the GluN1-N2A-N3A tri-NMDARs by measuring the COM distances of NTD-LBD, R1-R2, R2-D1, and D1-D2. Except for the D1-D2 distance, which was significantly longer in GluN1(C) compared to GluN1(A), the remaining length measurements and the vector angles between NTD and LBD were consistent between the two GluN1 subunits. We then extended the comparison to the GluN2A and GluN3A subunit in GluN1-N2A-N3A tri-receptors. The lengths of R1-R2 and R2-D1 in GluN3A were greater than those in GluN2A. We speculate that this loose domain arrangement contributes to the high instability of the GluN3A LBD. Additionally, the D1-D2 distance in the LBD of GluN3A was much shorter than that in GluN2A. These factors collectively result in a significantly larger vector angle between the NTD and LBD in the GluN3A subunit compared to GluN2A (Figure 2h).

### Activation mechanism of GluN1-N2A-N3A tri-NMDARs

Previous electrophysiological studies of GluN1-N2-N3A tri-NMDARs have relied on either co-expressing GluN1, GluN2, and GluN3A subunits in recombinant systems^53,54^ or recordings from native brain tissue^39,56^. However, in these experiments, GluN1-N2A- N3A tri-NMDARs may coexist with various other types of NMDARs, and the absence of selective pharmacological tools has hindered the biophysical characteristics of pure GluN1-N2-N3A tri-NMDARs. To address this limitation, we reconstituted purified GluN1- N2A-N3A tri-NMDARs into liposomes and performed single-channel recordings on these proteoliposomes.

To confirm that NMDARs form functional ion channels in liposomes, we initially reconstituted and recorded GluN1-N2A di-NMDARs with previous reports^64,65^. These GluN1-N2A di-NMDARs exhibited a super-cluster model of opening pattern, with an open probability of 0.46 ± 0.02, a mean dwell time of 17.07 ± 1.6 ms, and a unitary amplitude of 6.16 ± 0.12 pA (table 1). These parameters closely matched those biophysical properties recorded from HEK293 cells ^25^.

**Table 1.** Single-Channel Properties of N1-N2A, N1-N3A, and N1-N2A-N3A receptors.

| | N1-N2A<br>(100 μM Gly + 100 μM Glu) | N1-N3A<br>(3 μM Gly) | N1-N2A-N3A | |
| --- | --- | --- | --- | --- |
| | | | (3 μM Gly + 100 μM Glu) | (100 μM Gly + 100 μM Glu) |
| Open probability | $0.46 \pm 0.02^*$ | $0.05 \pm 0.01^*$ | $0.05 \pm 0.01^*$ | $0.17 \pm 0.01$ |
| Open probability 1 | $0.46 \pm 0.02$ | $0.05 \pm 0.01$ | $0.04 \pm 0.01$ | $0.04 \pm 0.01$ |
| Open probability 2 | / | / | $0.005 \pm 0.001^*$ | $0.07 \pm 0.01$ |
| Amplitude 1 (pA) | $6.16 \pm 0.12^*$ | $3.26 \pm 0.08$ | $3.63 \pm 0.04^*$ | $3.54 \pm 0.11$ |
| Amplitude 2 (pA) | / | / | $5.78 \pm 0.12^*$ | $5.93 \pm 0.04$ |
| Dwell time 1 (ms) | $17.07 \pm 1.6^*$ | $6.60 \pm 1.54^*$ | $2.25 \pm 0.37$ | $1.45 \pm 0.79$ |
| Dwell time 2 (ms) | / | / | $3.09 \pm 0.72^*$ | $10.52 \pm 3.56$ |
| Interevent interval 1 (ms) | $36.52 \pm 2.11$ | $153.10 \pm 40.72^*$ | $79.82 \pm 10.66$ | $46.90 \pm 7.76$ |
| Interevent interval 2 (ms) | / | / | $150.60 \pm 25.21^*$ | $237.80 \pm 44.08$ |

Values are mean ± SEM. \* $P < 0.05$ , unpaired t test, means significantly different from N1-N2A-N3A (100 μM Gly + 100 μM Glu)

We next recorded GluN1-N3A di-NMDARs with the glycine concentrations ranged from 0.1 μM to 100 μM (Extended Fig. 6, Extended table.1). Consistent with prior two-electrode voltage clamp recording^43,49^, we found that the open probability (0.052 ± 0.007) and dwell time (6.60 ± 1.54 ms) of GluN1-N3A di-NMDARs increased with glycine concentration, peaking at 3 μM. Beyond 10 μM glycine, the open probability declined, indicating that the reconstituted receptors retained the rapid desensitization characteristic of GluN1-N3A di-NMDARs.

To gain insight into the single-channel properties of GluN1-N2A-N3A tri-NMDARs and make comparisons with these of GluN1-N2A and GluN1-N3A di-NMDARs, we conducted recordings on reconstituted tri-NMDARs (Fig. 3a). We first examined whether the application of either glycine or glutamate alone could activate GluN1-N2A- N3A tri-NMDARs. No detectable currents were observed when either glycine or glutamate was applied individually (Fig. 3b), suggesting that the activation of GluN1- N2A-N3A tri-NMDARs requires the binding of both glycine and glutamate.

**Figure 3.**
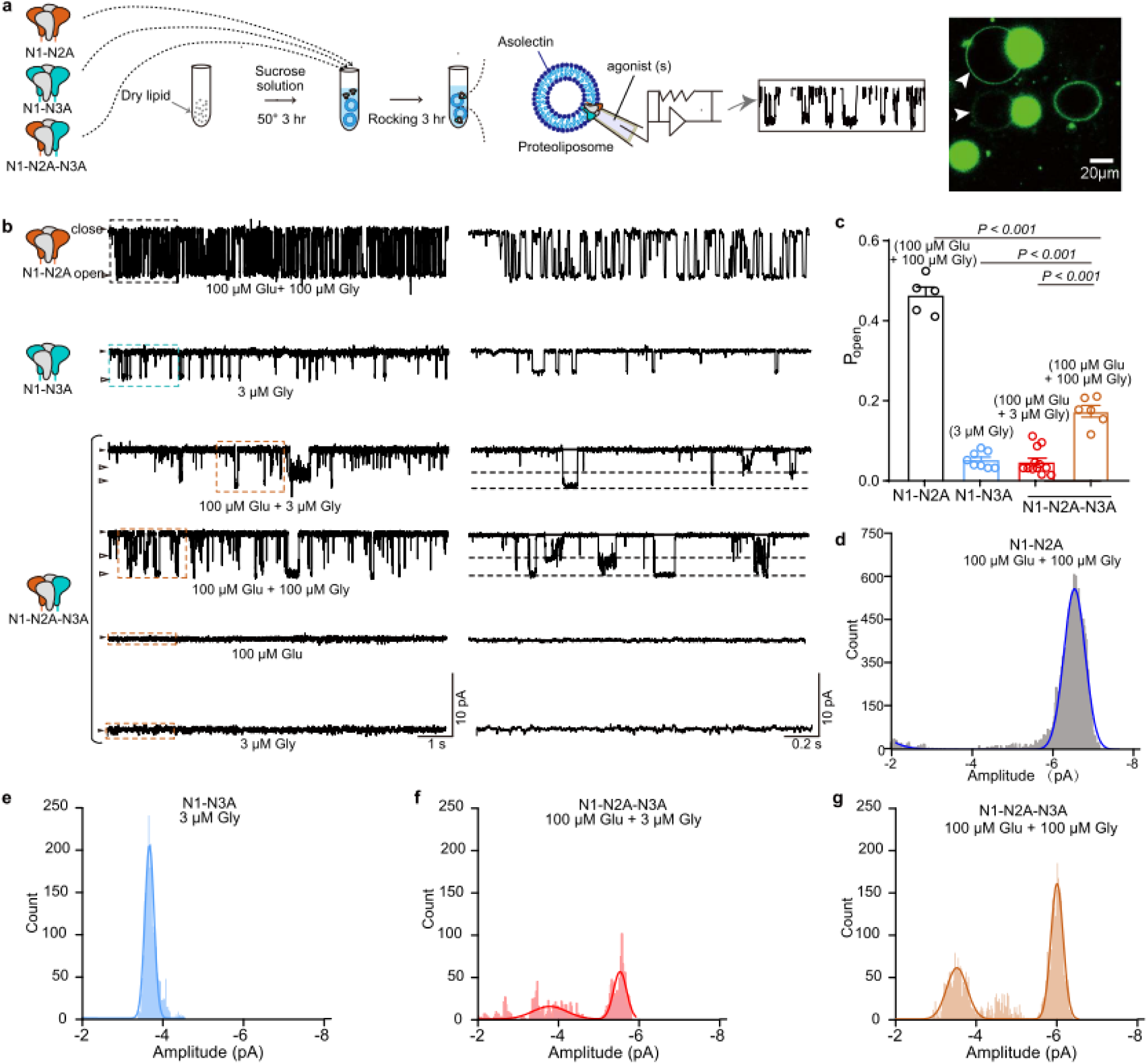
Single channel properties of proteoliposomes reconstituted with N1-N2A, N1-N3A, and N1-N2A-N3A receptors. **a**. Cartoons illustration of proteoliposomes reconstituted with N1-N2A, N1-N3A, and N1-N2A-N3A receptors, respectively. **b**. Representative single-channel currents of reconstituted N1-N2A, N1- N3A, and N1-N2A-N3A receptors at a holding potential of -60 mV with indicated agonist stimulation. Highlighted regions are shown on an expanded scale. **c**. Histogram showing single-channel open probabilities of N1-N2A (n=5), N1-3A (n=8), N1-N2A-N3A with 100 uM Glu plus 3 uM Gly treatment (n=12) and N1-N2A-N3A with 100 uM Glu plus 100 uM Gly treatment (n=6). The overall open probability values, from left to right, are 0.46 ± 0.020, 0.052 ± 0.007, 0.047 ± 0.010, and 0.172 ± 0.014. **d-g**. Amplitude histograms of single-channel conductance for N1-N2A, N1-N3A, and N1-N2A-N3A, fitted with the Sum of two Gaussian distributions.

Subsequently, we recorded the receptors under two glycine conditions 3 μM and 100 μM, in the present with fixed glutamate (100 μM). Unlike GluN1-N3A di- NMDARs, which desensitize at glycine concentration above 10 μM, GluN1-N2-N3A exhibited a higher open probability at high glycine (0.17 ± 0.01) compared to low glycine (0.05 ± 0.01), suggesting reduced glycine-induced desensitization. Also, unlike GluN1- N2A and GluN1-N3A di-NMDARs, which displayed only a simple unitary amplitude, the GluN1-N2A-N3A tri-NMDARs exhibited two distinct subconductance states in both glycine conditions. Specifically, under saturating co-agonist concentrations (100 μM glutamate and 100 μM glycine),, the GluN1-N2A-N3A tri-NMDARs showed two sublevels for the unitary currents (5.93 ± 0.04 pA and 3.54 ± 0.11 pA) with corresponding dwell times (10.52 ± 3.56 ms and 1.45 ± 0.79 ms). The open probability of GluN1-N2A-N3A tri-NMDARs (0.17 ± 0.01) was over 3-fold higher than that of GluN1- N3A activated by low glycine (0.05 ± 0.01), but significantly lower than that of GluN1- N2A (0.46 ± 0.02) receptors under the same saturating conditions (Fig. 3b-3g, Table 1).

Our findings demonstrate that the gate activation in GluN1-N2A-N3A tri-NMDARs requires the binding of both glycine and glutamate. However, the lack of rapid desensitization under high glycine concentrations suggests that the GluN3A’s role in GluN1-N2A-N3A tri-NMDAR gating differs from its role in GluN1-N3A di-NMDAR gating. In summary, these results reveal that the tri-NMDARs exhibit unique biophysical properties distinct from GluN1-N2A and GluN1-N3A di-NMDAR subtypes, highlighting their specialized functional characteristics.

### Mapping the subunit arrangement of GluN1-N2-N3A using click chemistry

To further validate the subunit arrangement of GluN1-N2-N3A tri-NMDARs and explore their presence in the brain, we employed click chemistry to site-specifically incorporate the photo-cross linker *p*-azido phenylalanine (AzF)^66,67^ into NMDARs subunits via genetic code expansion^68^. The azido group of AzF serves as the chemical handle for engineering photo-cross linking between proteins^69,70^, This approach offers several advantages for exploring protein-protein interactions. First, it requires only a single mutation for crosslinking, as AzF can form a convent band with any other nature amino acid. Second, UV-mediated crosslinking via the azido group occurs within a few angstroms. Last, the incorporation is biorthogonal with single-residue precision.

To implement the AzF system in our study, we first introduced AzF to the NTD of GluN2A subunit in the GluN1-N2A di-NMDAR, whose high-resolution structures have already been elucidated (PDB code: 7EOS) ^4^. We selected K220, located in the α6 helix of the GluN2A NTD R1 lobe, as the site for AzF incorporation. Utilizing an established genetic code expansion system in *Xenopus laevis* oocytes^66^ and HEK 293T cells^71^, we co-transfected plasmids encoding GluN1, GluN2A^K220AzF^, orthogonal suppressor tRNA (Yam), and AzF-tRNA synthetase (AzFRs+Yam), we confirmed the expression of GluN2A^K220AzF^ via fluorescence and western blot (Fig. 4a, b, Extended Data. Fig 7a, 7b). Notably, mRuby fused with GluN2A exhibited a typical membranal protein-like distribution, suggesting that AzF incorporation did not disrupt the trafficking and membrane insertion of GluN1-N2A receptors.

**Figure 4.**
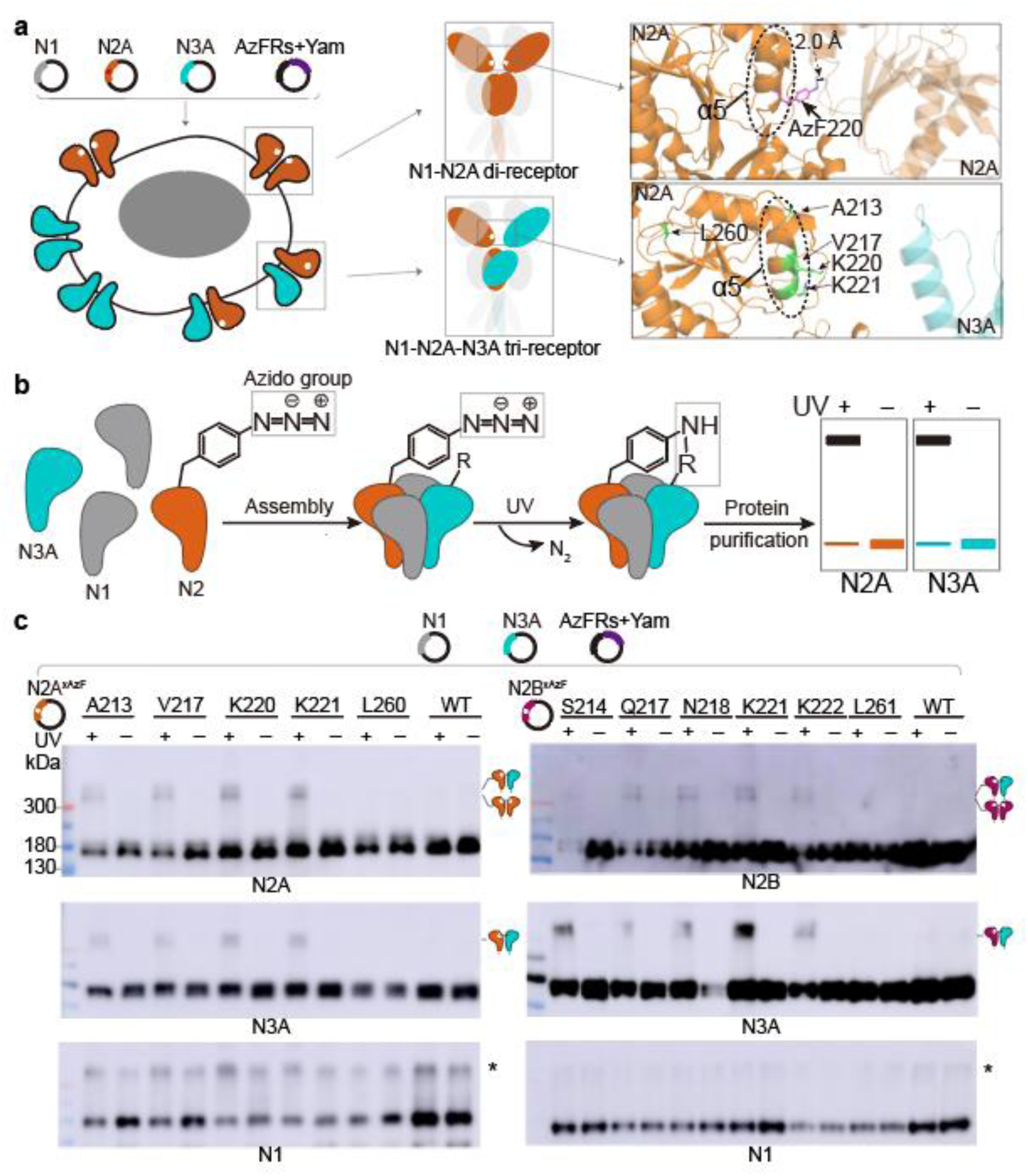
Probing the subunit geometry of N1-N2A-N3A and N1-N2B-N3A tri- receptors by click chemistry. **a**. Introduction of AzF in and N1-N2A di-NMDARs and N1-N2A-N3A tri-NMDARs through co-transfection of plasmids encoding N1, N2A with TAG mutant, N3A, engineered tRNA synthetase (AzFRs), and suppressor tRNA (Yam) (Left). Cartoon and model indicating AzF introduction sites at α5 of N2A NTD in N1-N2A and N1-N2A-N3A receptor (Middle and Right). **b**. Introduction of AzF at the interface of two NMDAR subunits. Upon UV stimulation, AzF-incorporated subunits form a covalent bond with nearby residues, detectable by western. **c**. Representative western blots demonstrating antibody recognition of N1, N2A or N2B, and N3A from cells under different treatments as indicated above the panels. Crosslinked protein bands are indicated by receptor Icon. A non-specific band was detected by anti-GluN1 as indicated by asterisks

Subsequently, we aimed to investigate whether the AzF-incorporated NMDARs could conduct current. In the presence of AzF, currents were recorded in nearly half of the injected oocytes (38 out of 78) after a 72-hour incubation with AzF. In contrast, no response was recorded in oocytes incubated without AzF, except in one oocyte with current over 100 nA (Extended Data. Fig 7C). Additionally, we examined whether UV illumination influenced NMDAR function. UV stimulation during agonist application increased the receptor current, indicating that a photo-switching effect mediated by AzF. To investigate whether this effect depended on photo-crosslinking, we conducted western blot analysis. After UV irradiation, a distinct band with a molecular weight exceeding 300 kDa, presumably corresponding to the crossed GluN2A-N2A dimer was resolved, confirming the UV-dependent GluN2A-N2A dimer formation (Extended Data. Fig. 7b-e).

Next, we co-expressed GluN2A^K220AzF^ with GluN1 and GluN2B to test whether AzF-incorporated GluN2A crosslink with another subunit. Western blot results showed a dimer with a molecular weight (over 300 kDa) detectable by both GluN2A and GluN2B specific antibodies, suggesting the formation of a UV-dependent GluN2A-N2B dimer. In summary, these results demonstrate that the incorporation of AzF into NMDARs and UV-induced capture of covalently crosslinked hetero-dimers (Extended Data. Fig. 7f).

Based on our cryo-EM structure of GluN1-N2A-N3A tri-NMDARs, we selected four sites (A213, V217, K220, K221) on GluN2A for AzF incorporation. These resides are located on the α5 helix of GluN2A-NTD, and their side chains likely face GluN3A-NTD . As a negative control, we chose L260, a buried residue within GluN2A-NTD (Fig. 4a).

When cells were irradiated with UV light, high-molecular-weight crosslinked bands were detected using antibodies against GluN2A and GluN3A, at these four α5 mutants (Fig. 4a, 4c).For the GluN1 subunit, only single monomer bands were shown up. This result suggests that GluN2A directly crosslinks with GluN3A at these sites. It is noteworthy that the anti-GluN2A antibody detected more than two high-molecular-weight bands, which may indicate the formation of crosslinked GluN2A homo-dimers. In contrast, cells expressing the negative L260 mutant and wildtype showed no detectable crosslinking bands.

Additionally, based on our GluN1-N2A-N3A tri-NMDARs structure, we constructed a homologous model of the GluN1-N2B-N3A tri-NMDARs. Using this model, we incorporated the AzF to five individual residue sites (S214, Q217, N218, K221, K222) located on the α5-helix of GluN2B-NTD, as well as a negative control site (L261) buried inside GluN2B-NTD. Consistent with the GluN1-N2A-N3A tri-NMDARs, crosslinking bands were detected by antibodies against GluN2B and GluN3A, but not GluN1 subunit (Fig. 4c), indicating that GluN2B and GluN3A are in a proximal position in the tetrameric NTD formation of GluN1-N2A-N3A tri-NMDARs. These results collectively validate our structural discovery that the NTD layer of GluN1-N2A-N3A tri-NMDARs adopt a GluN1- N2A-N1-N3A arrangement, with opposing GluN1 subunits and diagonally positioned GluN2A and GluN3A. To note, UV irradiation induced crosslinking in a small fraction of the dimers, demonstrating that only a subset of GluN1-N2-N3A tri-NMDARs meet the short spatial requirements for AzF crosslinking. This observation likely reflects the structural heterogeneity of GluN1-N2A-N3A tri-NMDARs observed in our structural analysis.

### Photo-crosslinking endogenous GluN3A subunit in cultured cortical neuron

To validate the presence of endogenous GluN3A and lay the foundation for subsequent biochemical experiments, we first performed immunofluorescence staining. Strong GluN3A signal were readily detected in the somatodendritic compartment of neurons (Extended Fig. 2), corroborating previous reports that GluN3A is endogenously expressed within these cellular domains^11,35^. Building upon this encouraging validation of endogenous GluN3A expression, we proceeded to exploit the potential for crosslinking between GluN3A and AzF-incorporated GluN2 subunits. To enhance the efficiency of AzF incorporation, we engineered a bicistronic plasmid with distinct cistrons. One encoded AzFRs/Yam, with AzFRs being accompanied by a red fluorescence protein and a flag tag, connected via a 2A oligopeptide sequence (AzFRs+Yam). The other cistron encoded GluN2A^K220AzF^, fused with green fluorescence protein and a strep tag. The transfection was confirmed through the detection of red fluorescence in plasmid-transfected groups, while green fluorescence was exclusively observed in the presence of AzF (+AzF), confirming AzF incorporation (Fig. 5a, 5b).

**Figure 5.**
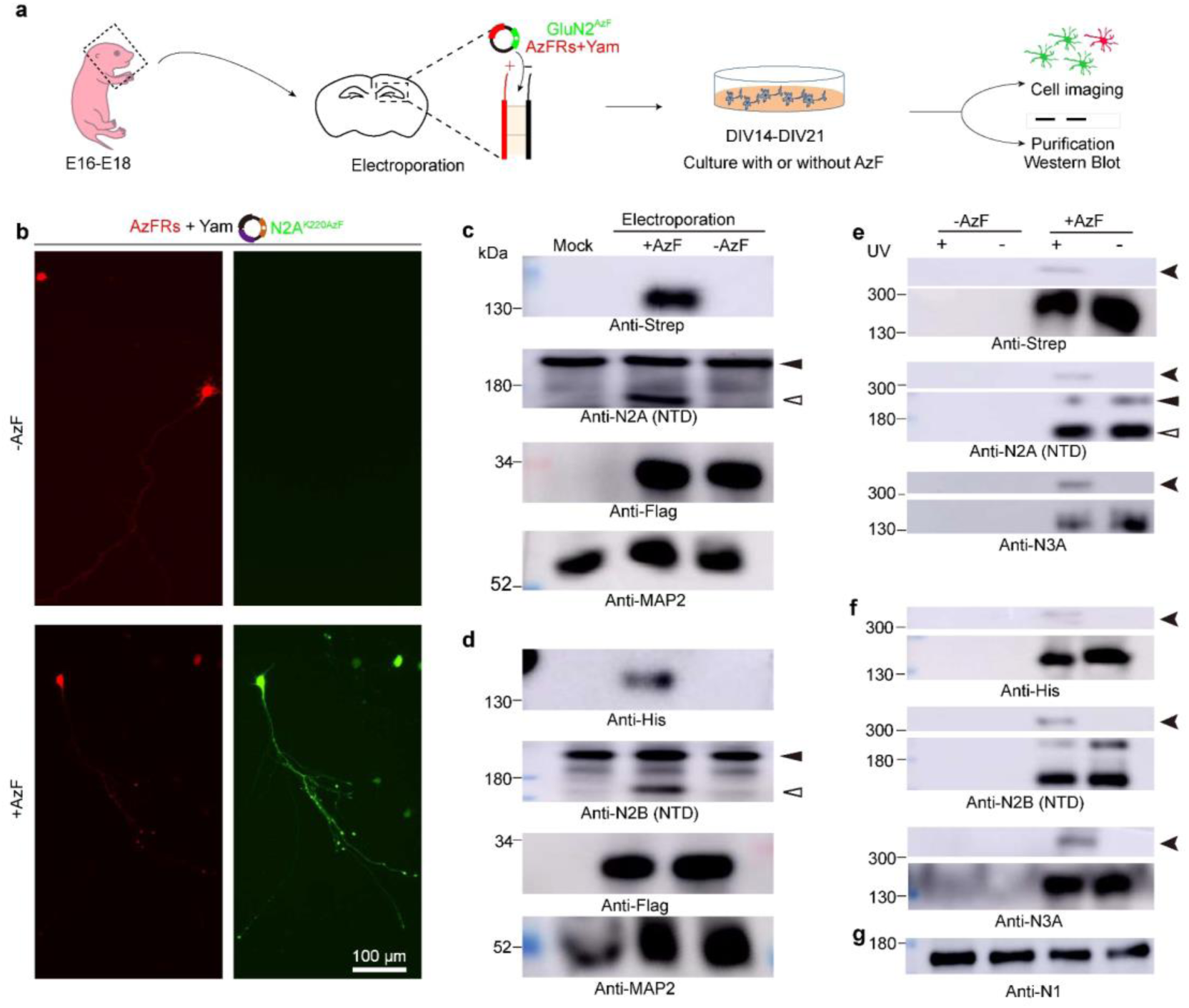
Proving the existence and the subunit geometry of N1-N2A-N3A and N1-N2B-N3A by applying click chemistry in cultured hippocampus neurons. a. **b.** Cultured neurons electroporated with bicistronic plasmids encoding engineered tRNA synthetase (AzFRs), suppressor tRNA (Yam), and N2A with K220AzF mutant. Electroporated neurons cultured with or without AzF. **c.** Representative western blots demonstrating antibody recognition of exogenously expressed GluN2A (Strep, c) or GluN2B (His, d), NTD of GluN2A or GluN2B, AzFRs (Flag), and MAP2 from cultured neurons electroporated with AzFRs-T2A-mRuby-Yam+hGluN2A^K220AzF^-GFP (c), AzFRs- T2A-mRuby-Yam+hGluN2B^K221AzF^-GFP (d) and cultured with or without AzF as indicated above the panels. Exaggerates GluN2A or GluN2B bands indicated by empty triangles, endogenous GluN2A or GluN2B by filled triangles. **e, f**. Representative blots for antibodies proved exogenous GluN2A (Strep, e) or GluN2B (His, f), NTD of GluN2A (e) or N2B (f), and GluN3A from Strep (e) or Ni-resin (f) purified examples extracted from the neurons electroporated with AzFRs-T2A-mRuby-4Yam+N2A^K220AzF^-GFP (e) or AzFRs-T2A-mRuby-Yam+hGluN2B^K221AzF^-GFP (f) and cultured with or without AzF as indicated above the panels. Crosslinked protein bands indicated by the arrowheads. **g**. Representative blots of GluN1 from 1% input for Strep purification.

Verification of AzF incorporation included the detection of AzF-incorporated GluN2 by antibodies targeting the N-terminal domain (NTD) of GluN2, along with the associated strep tags, resulting in a molecular weight of approximately 150 kDa. Notably, anti-GluN2A (NTD) antibodies exhibited reactivity toward both endogenous GluN2A (above 180 kDa) and exogenous AzF-incorporated GluN2A. The anti-Flag antibody effectively detected the Flag-fused AzFRs in all transfected groups (Fig. 5c).

After UV stimulation, AzF-incorporated GluN2A was isolated through strep-tag purification and subjected to western blot analysis. Antibodies directed against both GluN2A and GluN3A detected crosslinked bands exceeding 300 kDa following UV irradiation (Fig. 5e). These results affirm that AzF-incorporated GluN2A was instrumental in crosslinking with the endogenous GluN3A subunit. Using a comparable approach with K221AzF incorporated into GluN2B, we detected crosslinked GluN2B- N3A (Fig. 5d, 5g).

Taken together, the AzF incorporated GluN2A or GluN2B can covalently crosslinked with native GluN3A in cultured neuron, suggested that GluN1-N2-N3A tri- NMDARs assembled and may share a similar subunit establishment as the recombinant expressed receptors.

### Crosslinking of AzF-incorporated GluN2 with native GluN3A

Using the immunofluorescence staining, we proved that the GluN3A expressed in the cortex (Extended Fig. 2), as demonstrate by scRNA-seq (Fig. 1, Extended Fig. 1) and functional report^23,26^. To further investigate the potential crosslinking between AzF-incorporated GluN2 and native GluN3A subunit in the brain, we employed *in-utero* electroporation coupled with *in vivo* click chemistry. On embryonic days 14-15 (E14-15), we co-electroporated neonatal mice with constructs harboring AzFRs/Yam and GluN2A^K220AzF^. Post-electroporation, pregnant mice were provided with AzF containing water, and brains from juvenile mice were collected between postnatal days 7 to 14 (Fig. 6a), a period^18,21^ during which GluN3A subunit expression is known to be notably high.

**Figure 6.**
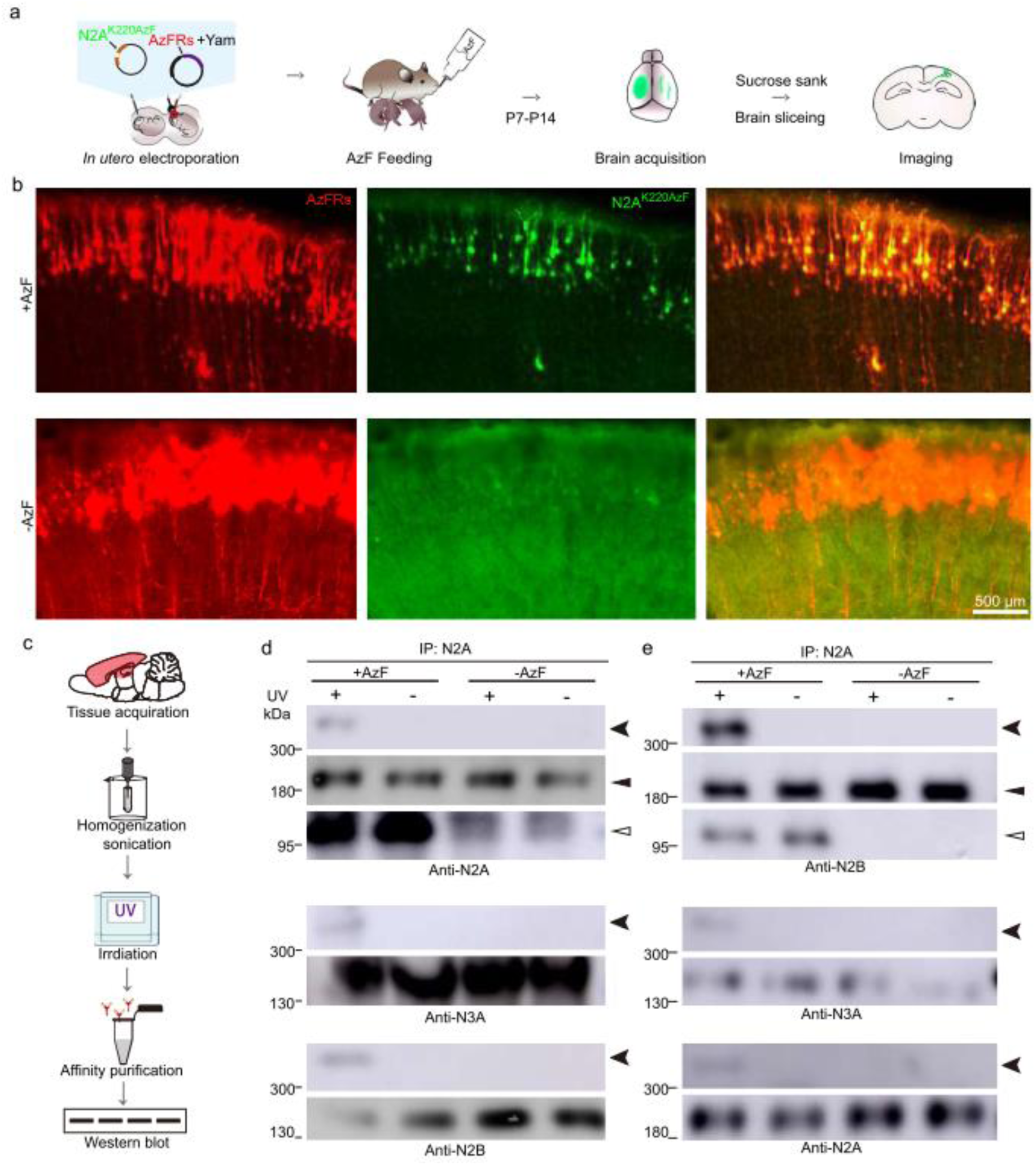
Proving the Existence and Subunit Geometry of N1-N2A-N3A and N1-N2B-N3A through *in vivo* application of Click Chemistry. **a**. Cartoon illustration depicting the *in vivo* application of click chemistry. **b**. Representative images of brain slices from in utero electroporated animal brains bred with or without AzF. The embryos underwent *in utero* electroporation with plasmids encoding the engineered tRNA synthetase (AzFRs), the suppressor tRNA (Yam), and GluN2A with the K220AzF mutant. After electroporating the DNA construct into a lateral ventricle, the embryos were gently inserted into the abdominal cavity. The pregnant mice were provided with AzF-containing water, and the brains were acquired at postnatal days 7-14. **c-e**. Total proteins from the cortical-hippocampal regions of AzF-incorporated animals or negative control animals were immunoprecipitated with anti-GluN2A (**d**) or anti-GluN2B (**e**). The immunoprecipitated were subsequently examined by Western blotting using anti- GluN2A, anti-GluN2B, and anti-GluN3A antibodies. The arrowhead, filled triangle, and open triangle indicate the bands corresponding to the crosslinked complex, endogenous GluN2A or GluN2B, and overexpressed GluN2A or GluN2B, respectively.

**Figure 7.**
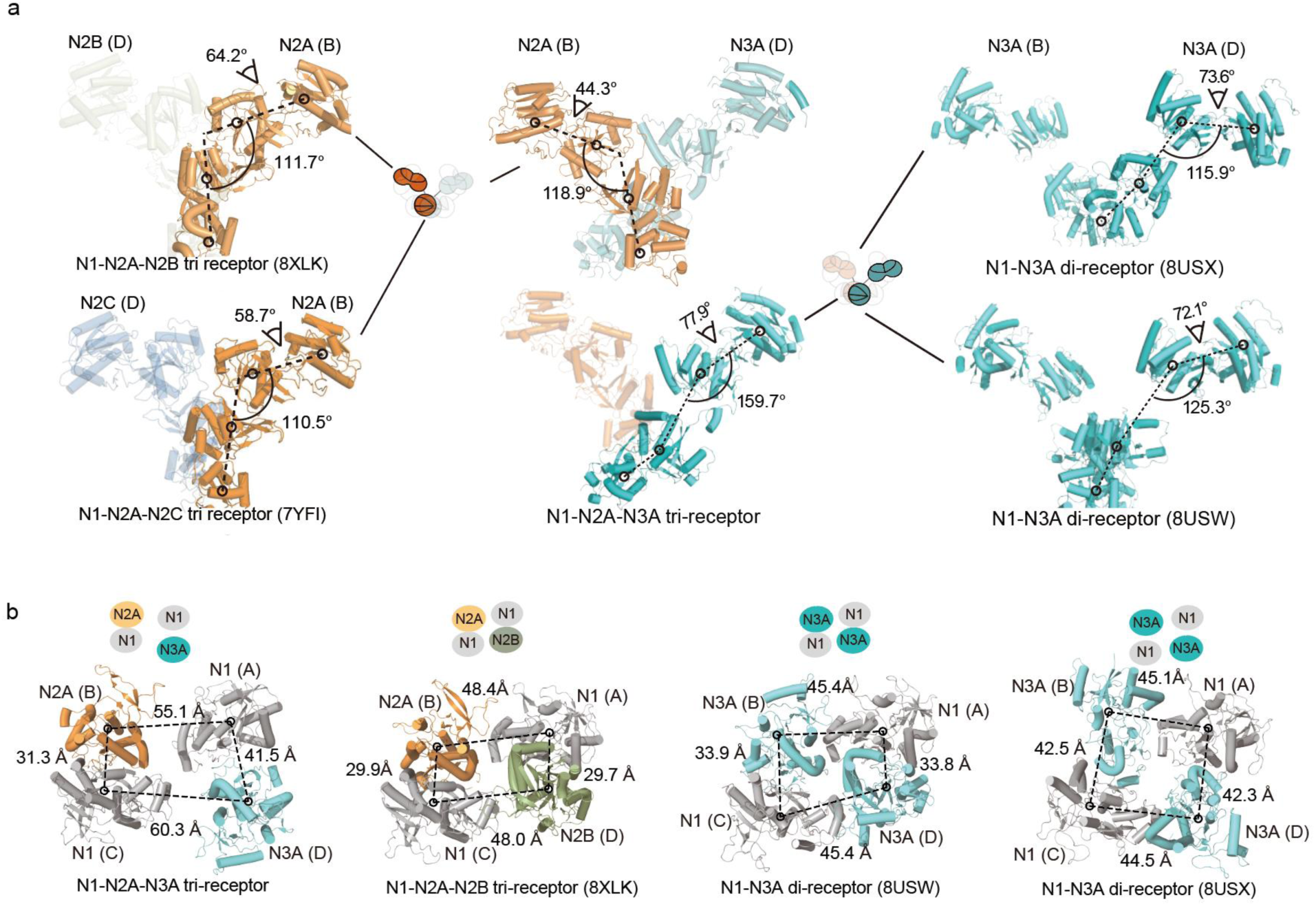
Structural comparation of N1-N2A-N3A with other NMDA receptors **a**. Structural analysis of extracellular domains of N2A or N3A subunits. The COM of R1 and R2 of NTD, D1 and D2 of LBD of each protomer is indicated by open circles. The COMs angles were measured. **b**. LBD comparison of N1-N2A-N3A with N1-N2A-N2B and N1-N3A.

The red fluorescence signal identified the location of electroporated neurons within the cortex region, where the GluN3A subunit has been reported highly expressed. In the with AzF groups, green fluorescence was observed within the red fluorescence-positive cells, confirming the incorporation of AzF into GluN2A (Fig. 6b). To further validate AzF incorporation, cortices from AzF animals were isolated, solubilized, and subjected to immunoprecipitation with an antibody targeting GluN2A (Fig 6c). Western blot results revealed that, in addition to the endogenous GluN2A (>180 kDa), a band with a molecular weight approximately 150 kDa was detected in the with AzF groups (Fig. 6d), confirming that the expression of AzF incorporated GluN2A.

To assess whether incorporated GluN2A could crosslink with other NMDAR subunits, brains from electroporated animals were homogenized and subjected to UV irradiation. Following enrichment by a GluN2A antibody, the samples were analyzed via western blot using antibodies specific to GluN2A, GluN2B, and GluN3A. Native GluN2A was detectable in all four experimental groups, indicating efficient immunoprecipitation.

Under UV stimulation, bands with high molecular weight were detected in the AzF-treated group, suggesting the formation of GluN2A^K220AzF^ complexes with other proteins. These bands were also detected by anti-GluN3A antibodies, providing evidence for the presence of GluN1-N2A-N3A complexes in the brain. Consistent with previous reports^72^, the existence of GluN1-N2A-N2B complexes was inferred, as high molecular weight bands were detectable with anti-GluN2B antibodies (Fig. 6d). Employing a similar strategy, we further demonstrated that AzF-incorporated GluN2B could form crosslinked complexes with GluN2A and GluN3A (Fig. 6e), thus confirming the presence of GluN1-N2B-N3A and GluN1-N2A-N2B receptors in the brain.

In summary, our findings provide compelling evidence for the existence of GluN1- N2-N3A tri-receptors in the brain through the *in vivo* application of AzF-incorporated GluN2, enabling the click-crosslinking of endogenous GluN3A subunits.

### Integrative summary of structural, electrophysiological, and in vivo findings on GluN1-N2A-N3A NMDARs

Together, our structural, electrophysiological, and in vivo experiments collectively establish GluN1-N2A-N3A as a distinct tri-NMDAR with unique subunit composition and functional properties. Structurally, the receptor adopts an asymmetric configuration at both the NTD and LBD layers, characterized by the presence of two GluN1 subunits, one GluN2A, and one GluN3A. Notably, the GluN3A subunit exhibits a highly flexible and loosely organized domain arrangement. This specific configuration at the NTD layer underlies its functional requirement for co-agonists glycine and glutamate, distinguishing it from the glycine-only activated GluN1-N3A di-NMDARs, and imparting unique single-channel characteristics that are not observed in GluN1-N2A or GluN1- N3A di-NMDARs.

Given the recent structural resolutions of multiple NMDARs, especially the native GluN1-N2A-N2B tri-NMDARs, we performed a comparative structural analysis focused on the alternative subunits. We examined conformational parameters such as domain clamshell angles and NTD-LBD bending. The GluN2A NTD of GluN1-N2A-N3A tri-NMDARs adopt a more closed conformation (R1-R2 clamshell angle: 44.3°) compared to its counterparts in GluN1-N2A-N2B (64.2°) and GluN1-N2A-N2C (58.7°), potentially due to steric effects from N2A-Fab_28C_ binding. The GluN3A displays a significantly more open and flexible conformation, with a larger R1-R2 angle (77.9°) and an exceptionally expanded NTD-LBD bending angle (159.7°), greater than that observed in di-NMDAR assemblies (115.9°, 125.3°)(Fig. 7a).

We then compared GluN1-N2A-N3A tri-NMDARs LBD layer with other NMDA receptors. Unlike other NMDARs, where domain-swapping in the NTD-LBD transduction results in the formation of two compact LBD heterodimers (A-D and B-C dimers), in GluN1-N2A-N3A tri-NMDARs, this transduction leads to only one compact heterodimer GluN1-N2A (A-D dimer) and one loose heterodimer GluN1-N3A (B-C dimer). The COM angle of the LBD between the A-D dimer is 17.5° larger than that between the B-C dimer, resulting in 10.2 Å shorter COM distance in the former compared to the latter (Fig. 7b). Notably, this asymmetry dimer-of-dimer LBD arrangement is not prominent in either endogenous GluN1-N2A-N2B tri-NMDARs (PDB 8XLK) or recombinant GluN1-N3A di-NMDARs (PDB 8USW, 8USX), as their COM length differences are less than 0.2 Å. Consequently, the LBD of GluN1-N2A-N3A tri- NMDARs is the most loosely organized, evidenced by the significantly larger total COM distances between the four subunits (> 13.8 Å), compared to the conventional GluN1-N2 di-NMDARs (Fig. 7b).

## Discussion

In this study, we provide direct evidence on the existence of GluN1-N2-N3A tri- NMDARs in the brain and reveal structural and biophysical properties of these receptors. By using scRNA-seq and immunoprecipitation, we found GluN3A subunits co-expressed and assembled with GluN2A and GluN2B subunits in various brain regions. By applying cryo-EM, we elucidated that GluN1-N2A-N3A tri-NMDARs possess an “GluN1-N2-N1-N3A” geometry and further validated the stoichiometry and *in vivo* existence of GluN1-N2A-N3A and GluN1-N2B-N3A tri-NMDARs by using click chemistry. Furthermore, we revealed biophysical properties of GluN1-N2A-N3A tri- NMDARs via proteoliposomes single channel recording. Our study enriched the understanding of sophisticated NMDAR subunits assembly and channel properties.

NMDA receptor family is composed of di-^24,73^ and tri^74–77^-subunit tetramers and with the tri-subunit tetramers are the dominant in the native brain^5^. Several GluN1-N2A di- NMDARs as well as tri-NMDARs, including GluN1-N2A-N2B^2,5,74,78^, GluN1-N2A- N2C^7,64,75^, and GluN1-N2B-N2D^76,79^, have been previously reported in the brain. These tri-NMDARs respond to a range of agonists, antagonists, and modulators specific to both their di-NMDARs counterparts. For example, GluN1-N2A-N2B tri-NMDARs can be modulated by the GluN1-N2A-specific positive allosteric modulator Zn^2+^[Ref^74^], as well as the GluN1-N2B-specific negative allosteric modulator ifenprodil^74^ and Ro 25-6981^2^. However, these tri-NMDARs exhibit distinct structural, pharmacological, and functional characteristics when compared to their di-NMDRs counterparts. Typically, tri-NMDARs display intermediate open probabilities and amplitude. The EC_50_ values for glutamate in

GluN1-N2B-N2D, GluN1-N2B, and GluN1-N2D receptors are approximately 0.5 μM, 0.27 μM, and 1.1 μM^76^, respectively. The main single-channel open amplitude for GluN1-N2A-N2C, GluN1-N2A, and GluN1-N2C are approximately 4.5 pA, 6.5 pA, and 3.5 pA^75^. In this study, we discovered that the gating of GluN1-N2A-N3A tri-NMDARs necessitates the presence of both glycine and glutamate, suggesting the independent role of GluN2A subunit and GluN1/GluN3A subunits in gating. Additionally, the incorporation of GluN2A renders GluN1-N2A-N3A tri-NMDARs less sensitivity to glycine compared to the GluN1-N3A di-NMDARs (Fig. 3b-g, Table 1, Extended Data Fig. 6, Extended Data Table 1).

GluN3A subunit, although discovered later^80^, have garnered increasing attention in recent years. Functional studies have demonstrated enhanced currents in GluN1-N3A di-NMDARs following preincubation with CGP-78608^51^ or L-689,560^48^, and further behavior evidence suggesting their significance in emotional control and locomotion^23,24^. However, the potential association of GluN3A subunits with GluN2 subunits to form functional GluN1-N2-N3A tri-NMDARs in the brain has remained elusive. A recent study showed no significant differences between wild-type and GluN2A knock-out animal when application of glycine followed CGP-78608. Based on these findings, the authors suggested a limited presence of GluN1-N2A-N3A tri- NMDARs in the brain^23^. Nonetheless, it’s essential to emphasize that the impact of the GluN2A subunit on the activation of GluN1-N2A-N3A tri-NMDARs remains unexplored. Consequently, excluding the existence of GluN1-N2A-N3A tri-NMDARs without glutamate application may be premature^23^. In this study, we establish that GluN3A tends to co-express with GluN2A and GluN2B subunits, forming functional GluN1-N2A-N3A and GluN1-N2B-N3A tri-NMDARs within the brain.

Moreover, our structural data unveiled that GluN1-N2A-N3A tri-NMDARs adopt a layered dimer-of-dimers arrangement, with GluN1 in A and C positions and GluN2A and GluN3A in B and D positions. This asymmetric NTD layer adopting a “super splayed open” conformation, with GluN3A’s NTD protrudes the furthest from the hypothetical center axis than the other subunits (Fig. 7a), which in consistence with the found that GluN3A is in a high flexibility in GluN1-N3A di-NMDARs^13^ .The most different of GluN1- N2-N3A tri-NMDARs to the conventional GluN1-N2A di-NMDARs^4^, GluN1-N2A-N2B^2^, or GluN1-N2A-N2C tri-NMDARs^64^ is the asymmetry conformation that characterized by the top-down view of NTD and LBD. Because the GluN1-N2A in GluN1-N2A-N3A remains relatively identical to GluN1-N2A NMDARs, we attribute this asymmetrical conformation from the incorporation of the GluN3A subunit. Unfortunately, our structure lacks sufficient TMD density and includes only glycine and glutamate-bound stage structures, precluding conclusions regarding LBD-TMD transitions and gating mechanisms.

To isolate GluN1-N2-N3A tri-NMDARs currents in a recombinant system is inherently challenging, given the co-expression of three subunits, resulting in a mixture of different receptor types. Furthermore, pharmacological tools to distinguish GluN1-N2- N3A tri-NMDARs from GluN1-N2 and GluN1-N3A di-NMDARs remain lacking. While the use of ectopic retention signals offers a delicate method for controlling subunit composition^74,76,78^, it may necessitate a systematic sequence for signal insertion, with improper insertion potentially yielding false results. To circumvent these challenges, we employed proteoliposomes-based single-channel recordings^81^ and detected GluN1- N2A-N3A currents in the presence of both glycine and glutamate. These results may service as an explanation for why GluN1-N2-N3A current is undetectable by preincubation of CGP-78608 following glycine^23^.

Our study also introduced click chemistry as a powerful tool to validate the geometry and *in vivo* existence of GluN1-N2A-N3A and GluN1-N2B-N3A tri-NMDARs. This was achieved by employing genetic code expansion to introduce the photo- crosslinker AzF (*p*-azido phenylalanine). The incorporation of AzF offers several advantages, including high consistency and specificity, as only a single mutation is required for comparison, and the crosslinking mediated by the azido group occurs within a few Å during UV stimulation. While prior studies have utilized photo-clickable unnatural amino acids (UAAs) for super-resolution imaging^82–84^ and ion channel manipulation in neurobiology^71^, our work represents the first application of photo- clickable UAAs to investigate direct hetero protein-protein interactions within the brain and underscores the potential of photo-clickable UAAs in neurobiology research.

In summary, our study provides compelling evidence for the existence of GluN1- N2A-N3A and GluN1-N2B-N3A tri-NMDARs through a combination of scRNA-seq, cryo-EM, single-channel recording, and photo-click chemistry. By addressing the long-standing controversy surrounding the existence of GluN1-N2-N3 tri-NMDARs in native systems, our research opens exciting avenues for further investigations into the structure and function of these receptors.

## Methods

### Generation and production of Fab-N1_4F11_, Fab-N2A_28C_, and Fab-N3A_1G4_

Fab-N1_4F11_, Fab-N2A_2-8C_, and Fab-N3A_1G4_ antibody fragments (fabs) were produced by papain digestion of N1_4F11_, N2A_28C_, and N3A_1G4_ monoclonal antibodies, respectively, and purified by cation-exchange chromatography as previously described^85^ with slight modifications. In brief, we used CTD-truncated human GluN1- N2A receptors (GluN1 residues 1-847, GenBank: NP_015566; GluN2A residues 1-841, GenBank: NP_000824) to raise N1_4F11_ and N2A_28C_ monoclonal antibodies and employed rat GluN1-N3A receptors (GluN1 residues 1–847, GenBank: NP_058706.1; GluN3A residues 1–965, GenBank: NP_612555.1) to raise the N3A_1G4_ antibody. These receptors were expressed in HEK293S GnTI^-^ cells (ATCC CRL-3022) through baculovirus transduction and purified using affinity and size exclusion chromatography (SEC) in TBS buffer (containing 20 mM Tris, 150 mM NaCl, pH 8.0), supplemented with 1% L-MNG, 2 mM Cholesteryl Hemisuccinate (CHS), 1 mM glutamate, and 1 mM glycine. Subsequently, these receptors were reconstituted into lipid-A-containing liposomes for immunization. The candidate antibodies were screened using immunocytochemistry and FSEC with GluN1-N2A, GluN1-N2B, GluN1-N2C, GluN1- N2D, GluN1-N3A, and GluN1-N3B receptors. The N1_4F11_, N2A_28C_, and N3A_1G4_ hybridoma cell lines produced antibodies that bound to GluN1, GluN2A, and GluN3A, respectively. To isolate the fabs, purified antibodies were cleaved by papain (Merck, 1495005) at a 1:30 w/w ratio in the presence of 5 mM cysteine and 1 mM EDTA for 2.5 hours at 37°C, followed by the addition of 30 mM iodoacetamide and incubation on ice for another 20 minutes in the dark. After removing the Fc fragment using protein-A agarose (Smart-life science), the free Fabs were further purified and concentrated by SEC in TBS buffer.

### Plasmid constructs

For cryo-EM studies, we used wild-type (WT) rat GluN1-1a (residues 1–847, GenBank: NP_058706.1), rat GluN2A (residues 1–847, GenBank: NP_036705.3), and rat GluN3A (residues 1–965, GenBank: NP_612555.1) constructs. To express and purify GluN1-N2A-N3A, the sequences encoding GluN1 and GluN2A were cloned into the pEG-BacMam Vector with a T2A ribosome skip sequence. The sequence encoding GluN3A was cloned into the pEG-BacMam vector. Both sequences contained an HRV 3C protease cut site, a four-alanine flexible linker, and an EGFP tag on the C-terminus, with a 6 X His-tag on GluN2A and a Strep-tag II on GluN3A, following EGFP. To improve expression levels and thermostability, a peptide fragment of the GluA2(Y837- K847) AMPA receptor was inserted after the M4 helix in GluN2A^86^. For AzF crosslinking experiments performed on HEK293S GnTI^-^ cells, we used pEG-BacMam vectors expressing wild-type human GluN1-1a, human GluN2A, human GluN2B (residues 1- 842, GenBank: NP_000825), and rat GluN3A with different AzF mutants, as described in the results section. The sequences encoded aminoacyl-tRNA synthetases (AzFRs) from *E. coli* TryRS, and the amber suppressor tRNA derived from *B. stearothermophilus* Tyr-tRNA_CUA_(Yam) was used as described previously^66^. The AzFRs sequence and Yam were promoted by CMV and U6, respectively. To improve AzF incorporation efficiency, we cloned AzFRs and four copies of Yam into the pEG-BacMam vectors with a poly(A) signal (AzFRs+Yam). For AzF crosslinking experiments in cultured neurons, we used the human GluN2A with the K220AzF mutation or GluN2B with K221AzF and AzFRs+Yam cloned into the bicistronic pEG-BacMam vector. In the AzF crosslinking experiment *in vivo*, we cloned the human GluN2A with the K220AzF mutation or GluN2B with K221AzF fused with a P2A ribosome skip sequence followed by an mNeonGreen tag into the CAGGS vector to improve electroporation efficiency. We cloned the AzFRs+Yam sequence fused with the T2A ribosome skip sequence, followed by an mRuby tag into the CAGGS vector. For TEVC recording, full-length human GluN1-1a, human GluN2A, and AzFRs+Yam were cloned into pcDNA3-based vectors. All site-directed mutagenesis was performed on the wild-type plasmid using Takara KOD-Fx DNA polymerase.

### Protein expression and purification

Protein expression and purification experiments were conducted as previously described^3,4,64^ with minor adjustments. Recombinant baculovirus containing the sequences encoding GluN1, GluN2A, and GluN3A were produced using sf9 insect cells following the instructions for the Bac-to-Bac TOPO Expression System (Invitrogen A11339). Suspended HEK293S GnTI^-^ cells at ∼3.5 × 10^6^ cells/ml were infected with P2 or P3 virus at a 1:1 molar ratio. Twelve hours after infection, 10 μM MK801 was added to prevent NMDAR-mediated excitotoxicity, and 10 mM sodium butyrate was added to boost expression. Cells were then incubated in a 30°C incubator for an additional 48-60 hours before harvesting. The following steps were performed at 4°C or on ice. Cells were resuspended in TBS with a protease inhibitor cocktail (1 mM PMSF, 0.8 mM aprotinin, 2 mM pepstatin, and 2 mg/mL leupeptin) and homogenized in a blender, followed by mild sonication for further disruption. An equal volume of lysis buffer (TBS with 2% L-MNG, 2 mM CHS) was added to the homogenate, and the samples were solubilized for 1.5 hours at 4°C. After ultracentrifugation at 40,000 rpm for 1.5 hours, the supernatant was collected and incubated with Strep-tactin beads for 1.5 hours. The protein was eluted with TBS buffer supplied with 0.1% L-MNG, 2 mM CHS, 5 mM D- desthiobiotin and further incubated with Ni-Resin supplied with 15 mM imidazole for 1 hour. Then the protein was eluted with TBS buffer supplied with 0.1% L-MNG, 2 mM CHS, and 200 mM imidazole. The eluted sample was incubated with 4F11 and 2-8C in a molar ratio of 10:1 and 5:1 for 30 min on ice. The sample was further purified by SEC using Superose increase 6 10/300 GL column in TBS buffer supplied with 0.1% digitonin, 5 μM CHS, 0.1 mM CHAPSO, 1 mM glycine, 1 mM glutamate, and 50 μM ethylenediamine tetraacetic acid (EDTA), at pH 8.0. The peak fraction was concentrated to 4–5 mg/ml. All purification procedures were conducted at 4 °C.

### Sample preparation and data collection

A droplet of 2.5 μl of 4 mg/ml fresh samples was applied to a glow-discharged Quantifoil grid 1.2/1.3 Au 300 mesh grids and then blotted and plunge-frozen in liquid ethane using FEI Vitrobot with 100% humidity at 8°C.The grids were subsequently screened by FEI Talos 120 V microscope and then collected on a 300 kV FEI Titan Krios cryo-electron microscope. The images were collected using a Gatan K3 direct electron detector in super-resolution mode at 29,000 x magnification (unbinned pixel size of 0.54Å /pixel). Images were collected using SerialEM or FEI EPU with a defocus range of -1.5∼ -2 μm. Each image stacks were recorded at 4 frames per second for 8 s, with a total dose is ∼60 e^-^/Å^2^.

### Image processing and model building

The movie alignment and particle picking were done using the program Relion 3.1^87^. Beam-induced motion and drift correction were performed using MotionCor2^88^.Defocus values were estimated using Gctf^89^. A total of ∼2000 particles were manually picked and subjected to an initial reference-free 2D classification. The 2D classifications with clear NMDAR features were used as templates for auto picking as implemented in the Relion. The auto-picked particles were subjected to rounds of 2D classification to clean up. The remaining were subjected to *ab-initio* 3D map generation, following rounds of 3D classification without imposing symmetry. After excluding the classes with incomplete density, the remaining classes were subjected to round 3D refinement and polishing.

Model building was done by docking individual lobes of NTD and LBD maps that were divided from 6MMP^1^, and predicted GluN3A^90^ to our structure using UCSF Chimera^91^. The PDB coordinates were manually inspected and corrected to fit into density using Coot 0.9.6.1^92^. The structure was further processed according to the density maps using Phenix (1.20.1)^93^ real space refinement with secondary structure and Ramachandran restraints. Local resolution of density maps is estimated by using ResMap 1.1.4^94^.

### Reconstitution of target proteins in Liposome

Reconstitution of the purified proteins into liposomes was conducted as described previously^81^. Briefly, the Asolectin (Sigma-Aldrich, 11145-50G) was dissolved in chloroform and dried under a stream of N_2_ in a glass test tube while the tube was rotated to make a homogeneous lipid film. After drying, a 5 μl of pure water was added to the bottom (pre-hydration) followed by 1 ml of 0.4 M sucrose. The solution was incubated for 3 hours at 50°C until the lipid was resuspended. After the solution cooled to room temperature, the purified proteins were shaken gently for an additional 3 hours at 4°C, then the samples were used for the patch-clamp experiments and observation under the confocal microscope.

### Patch-clamp and TEVC recording

The liposome patch-clamp recording was performed as previously reported protocol^81^. The pipette buffer was as follows (in mM, pH 7.2): 140 CsCl, 5 EGTA, 10 HEPES, and the bath buffer contained as follows (in mM, pH 7.3) 140 NaCl, 2.8 KCl,

0.5 CaCl_2_, 10 HEPES and 0.01 BAPTA. The glass pipettes (BF150-86-10, Sutter Instrument) used for recording were pulled using P-97 puller (Sutter Instrument) with resistance in the range of 3-6 MΩ.

The pipette formed a tight seal with the liposome membrane with a resistance over a Gigaohm seal (>1GΩ). The membrane was then pathed at -60 mV, the agonists were added to the pipette solution to a certain concentration and the channel opening was recorded. For the GluN1-N2A, we used 100 μM Glu and 100 μM Gly, for the GluN1-N3A, we used 3 μM Gly, and for the GluN1-N2A-N3A, we used 100μM Glu and 100μM Gly. All experiments were performed at room temperature, and all the recording data were collected with EPC-10 amplifier and Pulse software (HEKA Electronic, Lambrecht, Germany) with a 0.5-kHz low-pass filter and 50-Hz notch filter. Independent recordings of 10 s duration were used for data analysis.

*Xenopus laevis* oocytes were prepared and injected as described previously^66^. The constructs in pcDNA3-based vectors were nuclear injected into *Xenopus laevis* oocytes with 36 nl of a mixture of cDNAs as follows, GluN1 (50 ng/μl), GluN2A^k220AzF^(50 ng/μl), AzFRs+Yam (10 ng/μl) and after about 12 hr of incubation at 18°C in the Barth solution supplemented with gentamicin sulfate (50 ng/ml) and D-APV (50 μM). AzF was dissolved with sonication in Barth solution (stock solution at 10 mM) and diluted (2 mM) for oocyte incubation. Currents were measured using a two-electrode voltage clamp at a holding voltage of -60 mV. The stand external solution was (in mM pH 7.3): 100 NaCl, 5 HEPES, 2.5 KOH. NMDAR-mediated currents were induced by the simultaneous application of Glu and Gly at the concentration of 100 μM each. The heavy-metal chelator diethylenetriamine-pentaacetic acid (DTPA, 10 µM) was supplemented to all bath solutions.

The single-channel conductance was obtained through the linear fitting of the current-voltage plots. For quantitative analysis, the Pulse files were converted into PCLAMP format using the ABF file Utility. Current trace analyses and fits were done with Clampfit software. The oocyte data was collected and analyzed using pClamp 10 (Molecular Devices). The Statistical Program for Social Science (SPSS) and Graph Pad Prism 7 was used for statistical analysis and graph generation. The *t*-test and one-way ANOVA were used to assess statistical significance. The data are shown as the mean value ± s.e.m.

### Hippocampal neuron culture and electroporation

The hippocampal neurons were cultured as previously described^95^ with a small adjustment. In brief, the hippocampus was obtained from the 16-18-day-old Sprague- Dawley rat embryos. After removal of blood vessels, pia mater, and cortex, the freshly dissected hippocampus was sectioned into small fragments and digested with 0.25% trypsin for 15-30 min at 37 °C, and then terminated by the addition of DMEM with 10% fetal bovine serum. Then the treated tissue was dispersed mechanically. Dissociated neurons were transfected by electroporation using the Amaxa Necleofector device. Then and cells were plated at a density of 5 x 10^5^cells/ml on poly-L-lysine-coated dishes in DMEM with 10% fetal bovine serum. The medium was changed to neurobasal containing 2% B27 supplement the next day supplied with or without AzF. Then culture medium was changed twice a week. The 14–21-day-old neurons were used. The fluorescence detect experiment were performed as previously described with small adjustment^96^, the cells were imaged by a Nikon Ti2 microscope with the FITC and mRuby optical filter.

### *In-utero* electroporation and brain acquisition

The *in-utero* electroporation was performed on the 15-16-day-old ICR rat embryos^97^. Briefly, the pregnant mouse was anesthesia in the induction chamber with air flow at 1 l/min and isoflurane at 5% until the animal is sedated and continue treated with air flow at 1 l/min and isoflurane at 2%. Then the uterine horn was exposed, and the plasmids were injected into the lateral ventricle. The electroporation was performed by ECM830 Electroporator with 45V voltage, 2 pulses, 50 ms pulse duration, and 500 ms interval. After electroporated, the uterine horn was put back in the abdomen. For the AzF group, the AzF-containing drink was provided. The newborn mice were used on 14- day post birth. The brain section for the imaging was prepared as previously described^98^. Mice were anesthetized and transcranial perfused with 0.9% saline solution followed by a 4% paraformaldehyde dissolved in 0.1 M phosphate buffer (pH 7.4). The brains were removed and cryoprotected using a graded sucrose series (20%, 30%) until they sank. A freezing microtome was used to cut 30-60 μm sections for imaging. Brain slices were washed and then imaged by the VS120 microscope at excitation and emission wavelengths of 488 and 519 nm (mNeonGreen) and 591 and 618 nm (mRuby). The brains for immunoprecipitation and western blotting were obtained by transcranial perfused mice with 0.9% saline solution. Only the cortex and hippocampus were used for analysis

### Immunoprecipitation and Western blotting

The immunoprecipitation was performed as previously described^96^. The brain tissue was homogenized in a blender for 2-3 min and further disrupted by sonication at 4°C in TBS containing protease inhibitors. The homogenate was next subjected to solubilized and centrifuging. The supernatant was mixed with antibodies. For immunoprecipitation, proteins were incubated with antibodies at 4°C for 2 hours. Then incubated the protein-antibody complex substances with 100 μl protein G Sepharose CL-4B beads at 4°C for another 2 hours. Beads were then washed and eluted for western blotting. For the western blot, an equal volume of samples from each group was electrophoresed on 8% or 10% SDS-PAGE gels and then transferred to polyvinyl difluoride (PVDF) membranes. After being blocked by 10% milk in TBST at room temperature for 2 hours, the membrane was incubated with anti-Strep, anti-Flag, anti- GluN1, anti-GluN2A, anti-GluN2B, or anti-GluN3A at 4°C overnight. After washing in TBST, membranes were incubated with HRP-conjugated anti-rabbit IgG or anti-mouse IgG for 1 hour at room temperature. Then peroxidase activity was visualized by AI600.

### scRNA-seq analysis

scRNA sequence analysis was done using the publicly available data set^60–63^ and using previously described approaches^99^. Data was analysis using R Studio and Prism.

## Data availability

The cryo-EM density maps and corresponding coordinates have been deposited into the Electron Microscopy Data Bank and Protein Date Bank with the following accession codes: EMD-36209 and 8JF7 for the fabs bound state. The scRNA-seq database used in this study is available in the Gene Expression Omnibus of NCBI with the following accession codes: GSE185862, GSE121891, GSE146692, GSE165371.

## Acknowledgement

We thank Dr. Yu Kong and Lijun Pan at Electron Microscopy Facilities of Center for Excellence in Brain Science and Technology, Chinese Academy of Sciences for assistance with sample screen. This work was supported by National Natural Science

Foundation of China (NSFC) (32100762, to Z.K.)

## Author contributions

Z.K. conceived the project. C.X. and N.S. prepared subunit specific antibodies. Y.S. and B.W. analyzed the single-cell sequencing data. Z.K. and T.Z. froze the protein. F.Y. performed the proteolipsomes reconstruction and recording. Z.K. collected and analyzed cryo-EM data, built atomic models, performed gene construction, protein expression and purification, hippocampal neuron culture, *in-utero* electroporation, immunoprecipitation, western blotting, imaging, and TEVC recording. Z.K. designed the experiments and wrote the manuscript. All authors read and approved the contents of the manuscript.

## Declare of interest

The authors declare no competing interests.

---

**Extended Data Figure 1.**
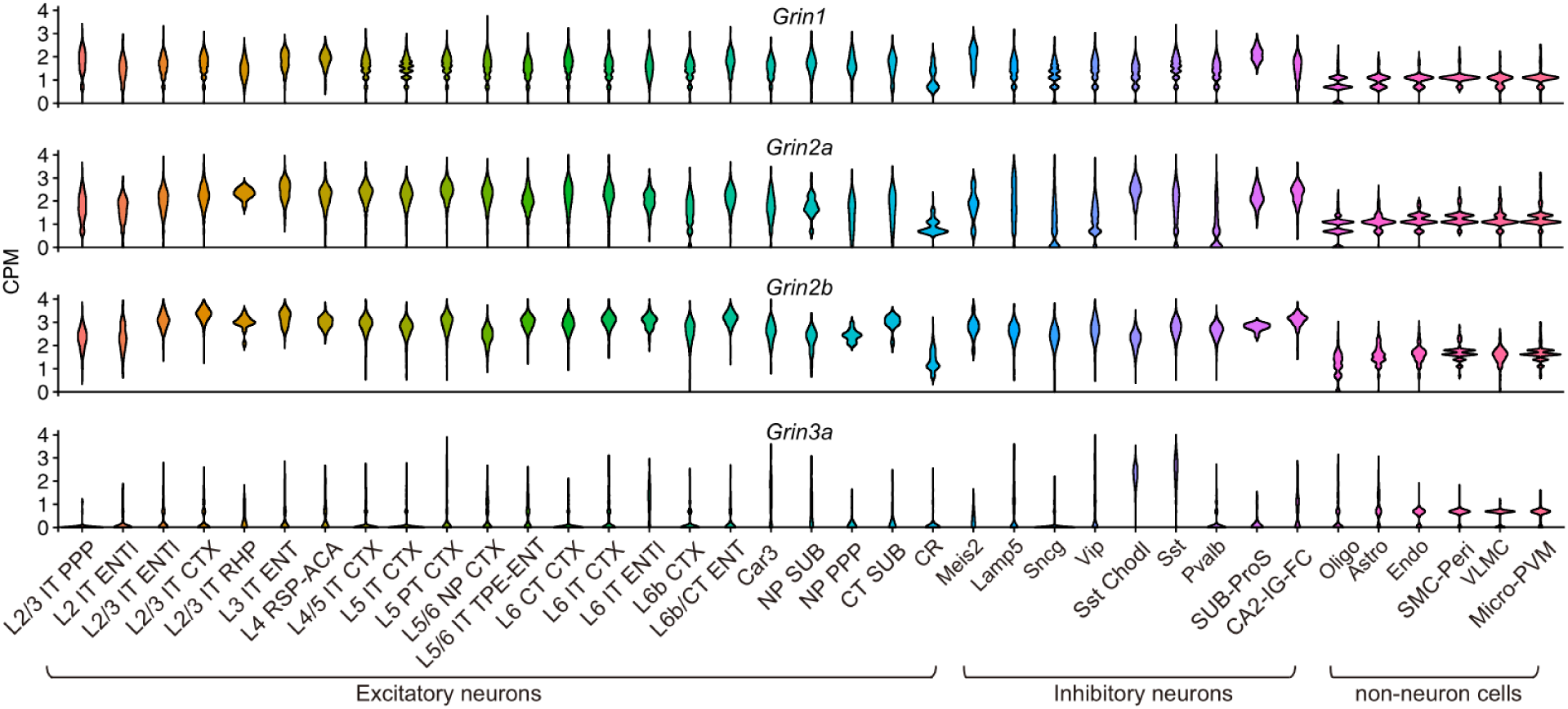
Violins plot showing the expression level of *Grin1*, *Grin2a*, *Grin2b*, and *Grin3a* from different cell types in mouse cortex and hippocampal formation.

**Extended Data Figure 2.**
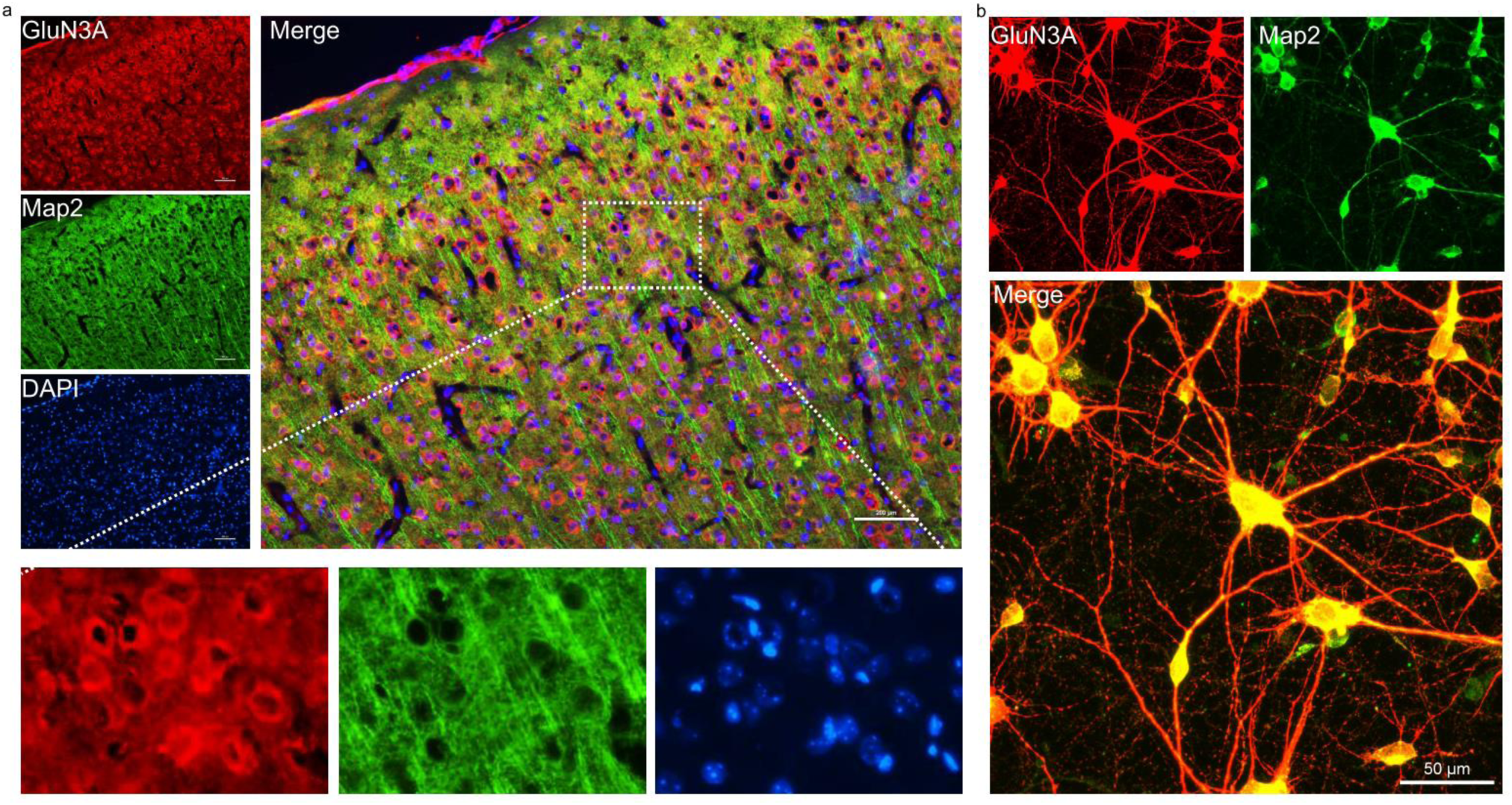
Representative immunofluorescence image of brain slice (a) and cultured hippocampal neurons (b) stained of GluN3A (red) and MAP2 (green).

**Extended Data Figure 3.**
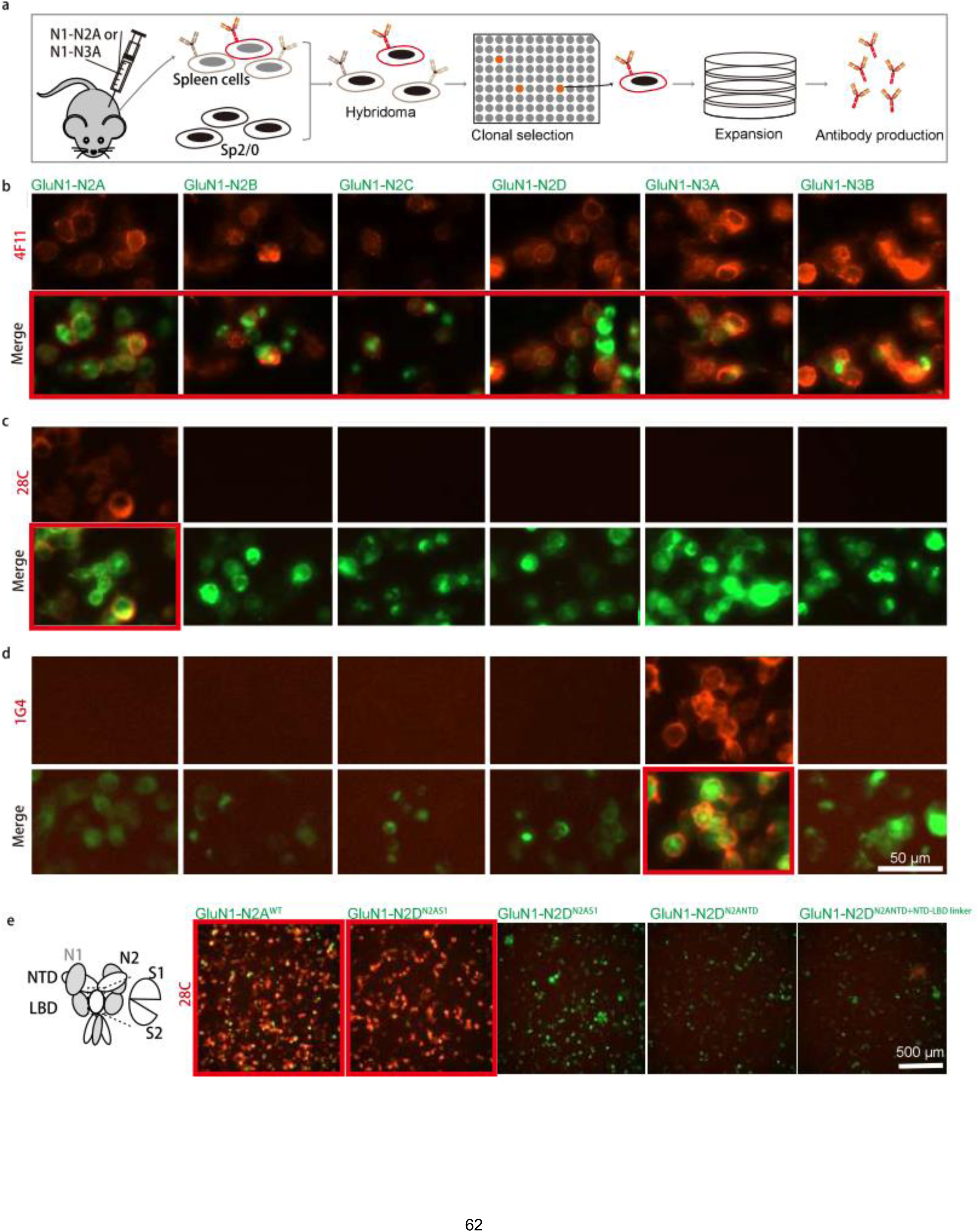
Generation and validation of NMDAR subunit-specific antibodies. a. Cartton illustration outlining the antibodies generation process. Full-length GluN1-N2A receptors were used for the development of anti-GluN1 and anti-GluN2A antibodies, while full-length GluN1-N3A receptors were employed for the development of anti-GluN3A antibodies. b-d. Representative immunofluorescent staining of HEK GnTI^-^ cells expressing GFP-tagged different subtypes of NMDARs using anti-GluN1 (clone No. 4F11, b), anti-GluN2A (clone No. 28C, c), and anti-GluN3A (clone No. 1G4, d) antibodies. e. Representative immunofluorescent staining of HEK GnTI^-^ cells expression GFP-tagged wild-type GluN1-N2A, wild-type GluN1-N2D, and various chimeric GluN1-N2D receptors.

**Extended Data Figure 4.**
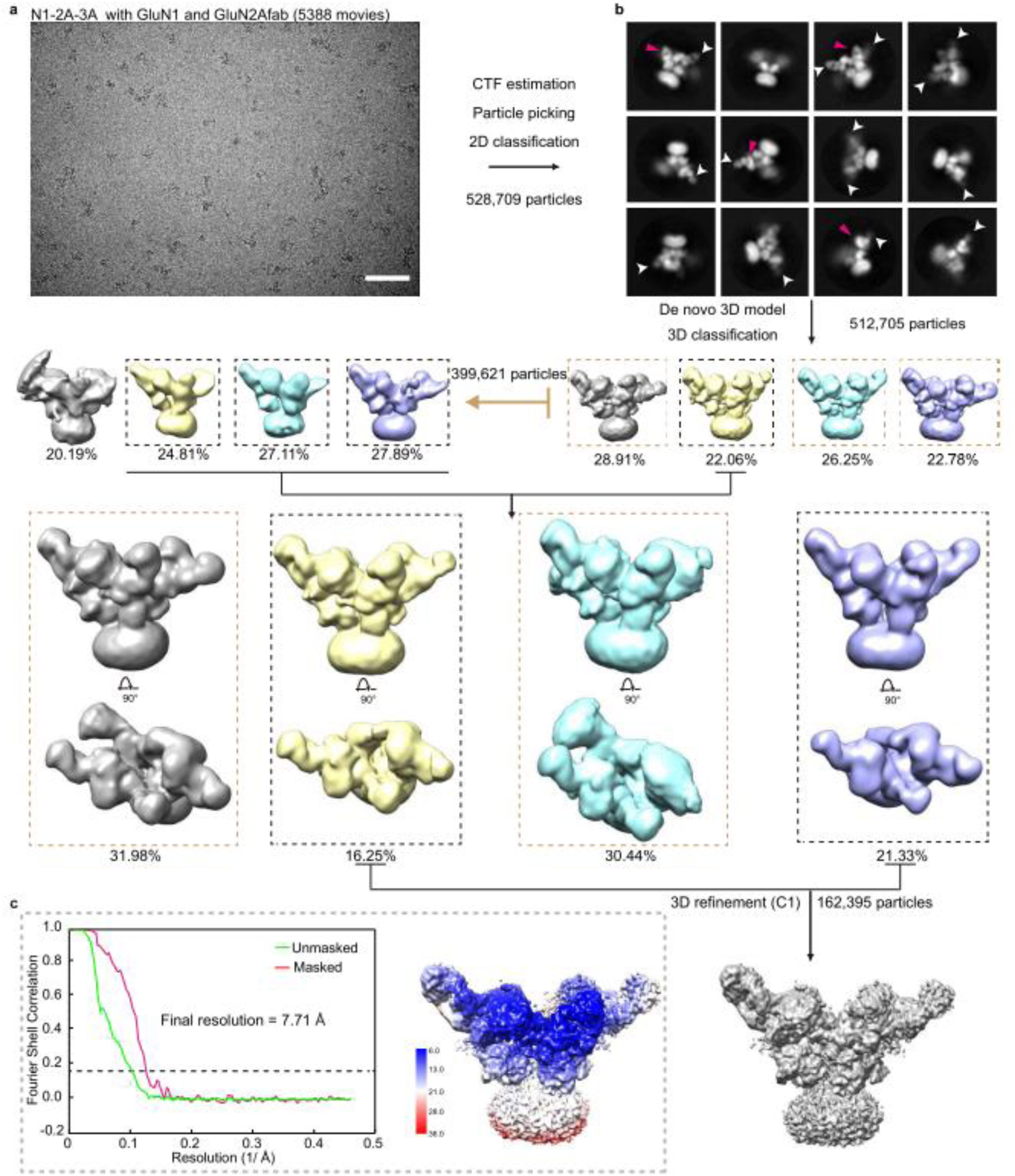
Overview of cryo-EM image processing and 3D reconstruction of Fabs bound GluN1-N2A-N3A receptor structures. a, b, Flowchart of image processing and 3D reconstruction of Fabs bound GluN1-N2A-N3A receptors. The 2D class images show Fabs bound 2D views in different orientations. The 3D classes with well features were selected and combined through several rounds of 3D classification for final refinement. In panel b, the red and white arrowheads represent localization of the anti- GluN1 fab or anti-GluN2A fab, respectively. c, Fourier shell correlation (FSC) curve for the resolution estimation.

**Extended Data Figure 5.**
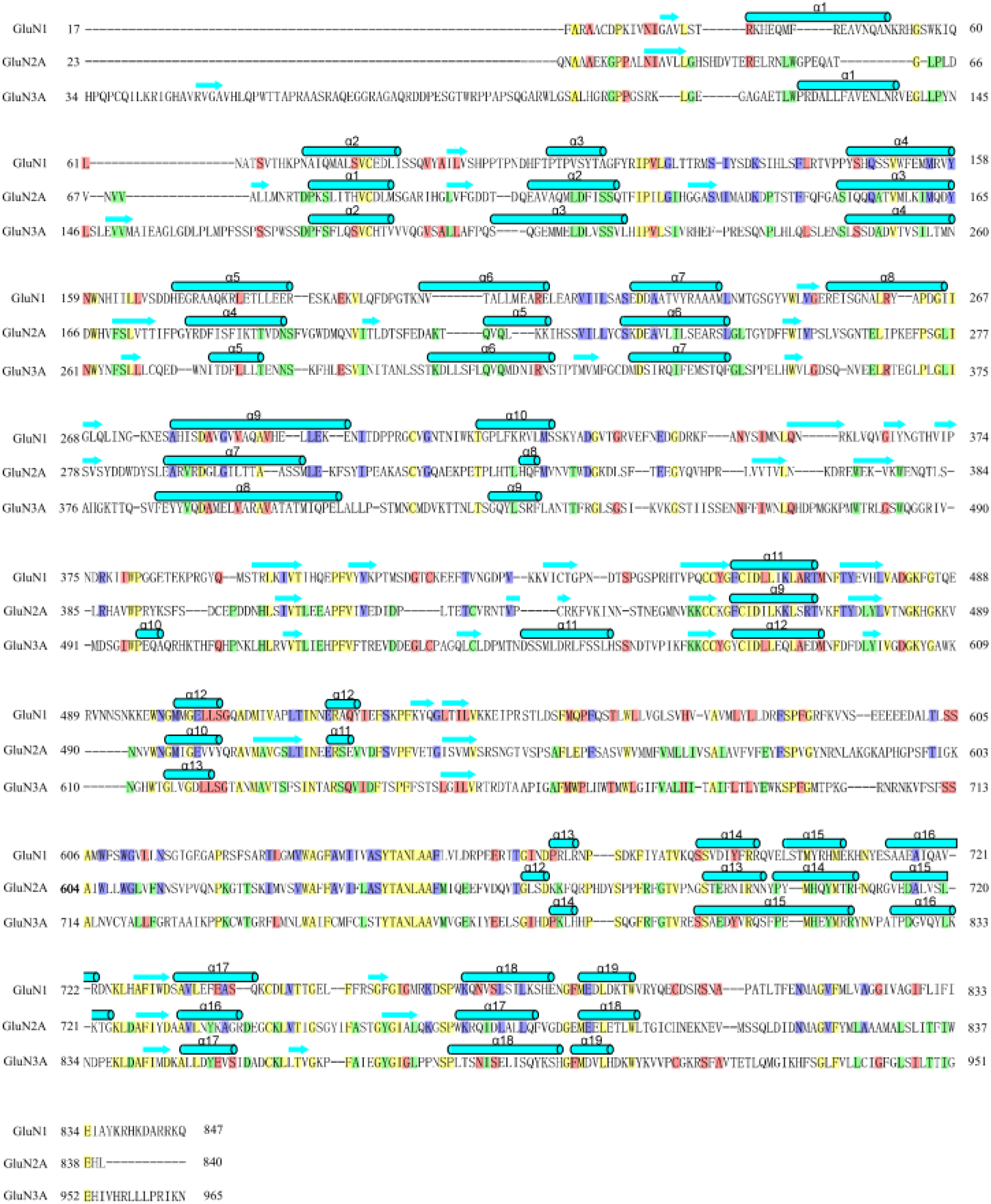
Sequence alignment of rat GluN1, GluN2A, and GluN3A. Secondary structure elements of GluN1, GluN2A, and GluN3A are indicated in above sequence alignment. Invariant hand highly conserved amino acids are sharded yellow. Green GluN2A vs GluN3A, pink (GluN1 vs GluN3A), and violet (GluN1 vs GluN2A).

**Extended Data Figure 6.**
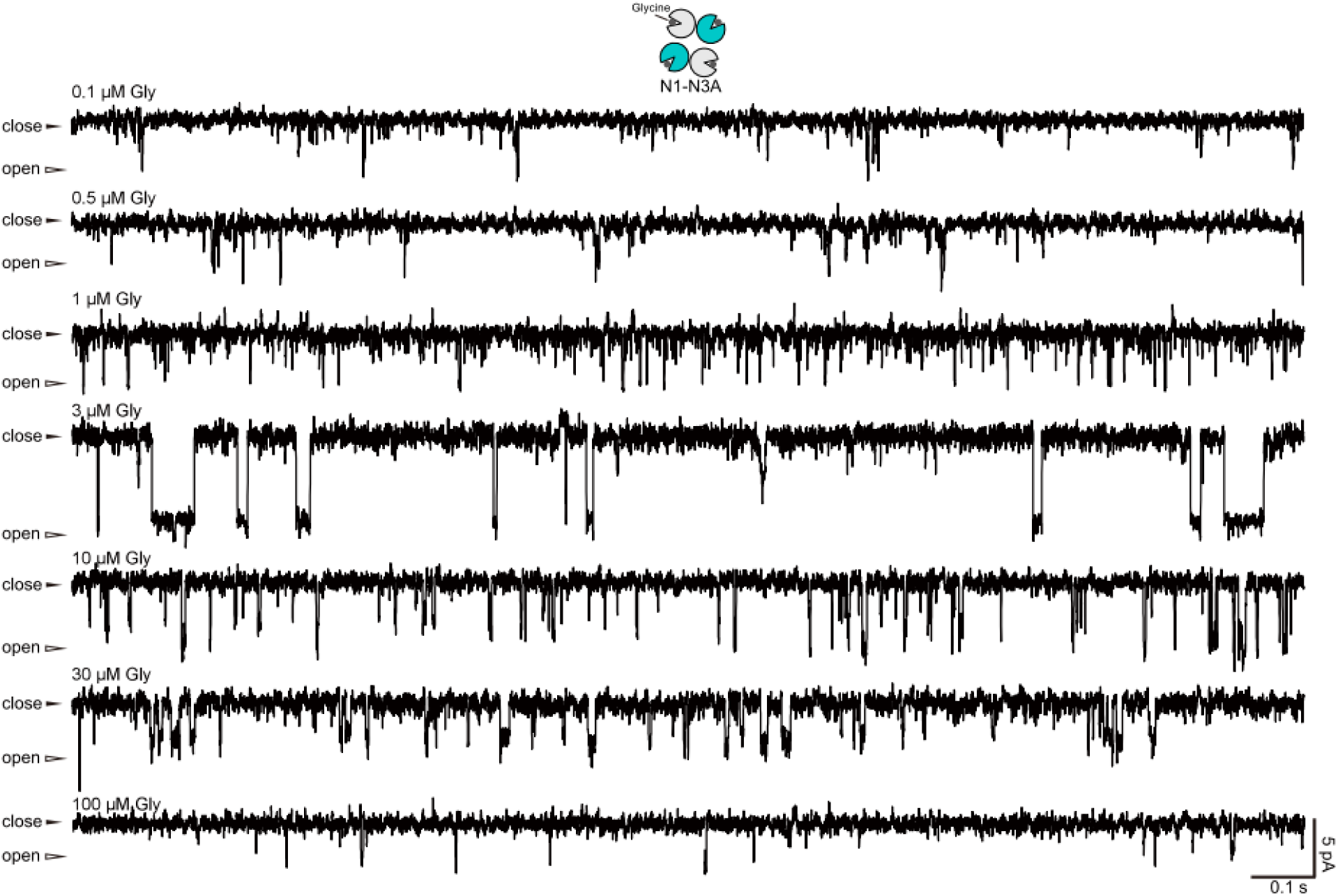
Channel currents from proteoliposomes reconstituted GluN1-N3A receptors with different concentration of Gly treatment .

**Extended Data Figure 7.**
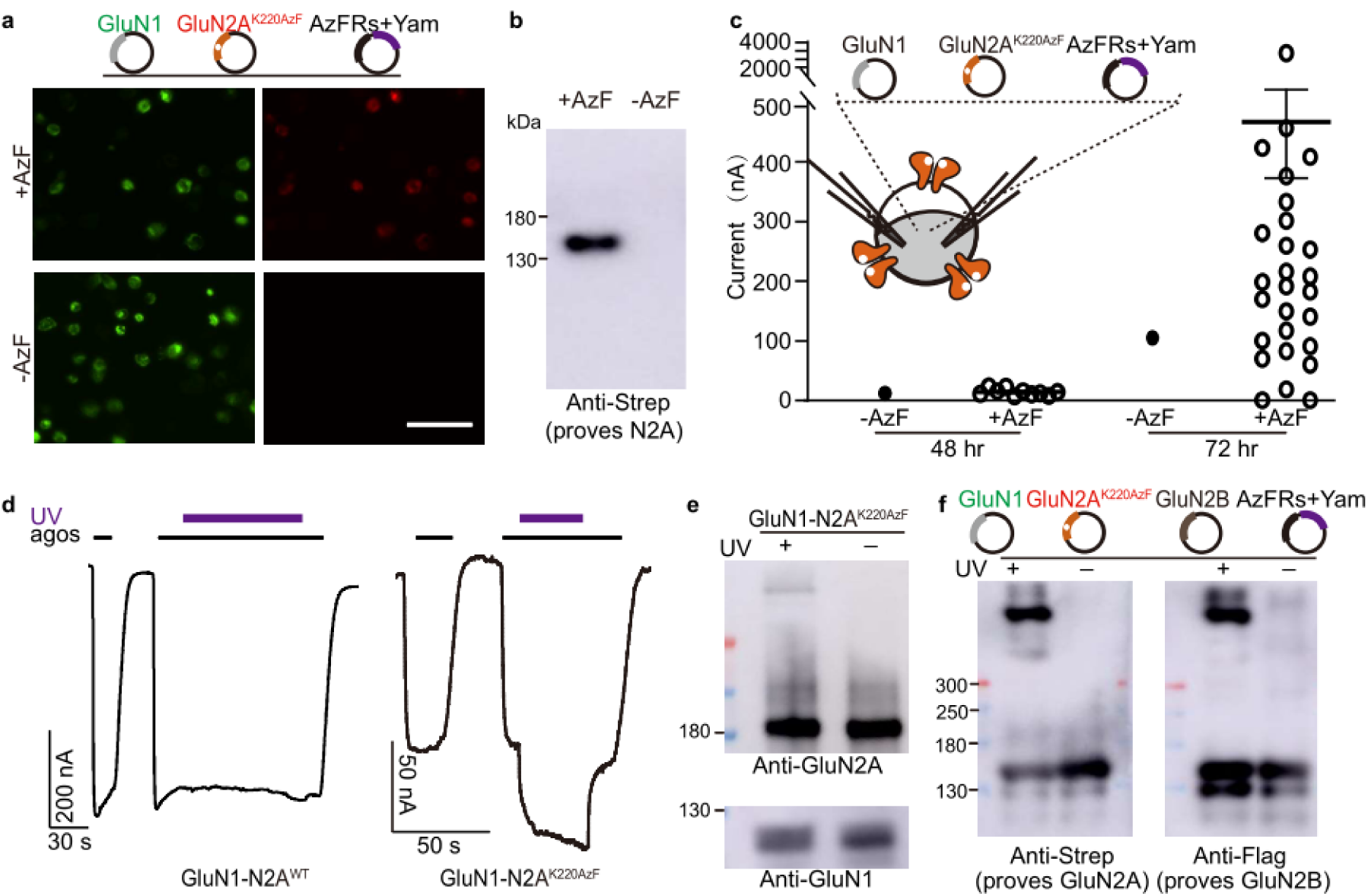
Incorporation of the AzF into GluN2A through genetic code expansion. a, b. Three plasmids encoding GluN1, and N2A with TAG mutant at the position of K220, and the engineered tRNA synthetase (AzFRs) and the suppressor tRNA (Yam) were transfected with HKE cells. Representative fluorescence images (a) and western blot (b) indicate the the incorporation of AzF into GluN2A. c. Three plasmids encoding N1, and N2A with TAG mutant at the position of K220, and the engineered tRNA synthetase (AzFRs) and the suppressor tRNA (Yam) were co-injected into Xenopus laevis oocytes. Injected oocytes were then cultured with or without AzF, and the current was recorded at 48 hr and 72 hr after injection. For each condition, more than 20 oocytes were tested. Only currents over 10 nA were plotted. d. Representative traces recorded on oocytes injected with GluN1-N2A WT and GluN1-N2A K220AzF and UV stimulation. e. Representative blot for antibodies against GluN1 and GluN2A from oocytes injected GluN1- 2A K220AzF with or without UV stimulation. f. Four plasmids encoding GluN1, GluN2A with TAG mutant at the position of K220, the engineered tRNA synthetase (AzFRs), and the suppressor tRNA (Yam), GluN2B were transfected with HEK cells. Transfected cells were cultured with AzF, and the AzF-incorporated GluN1-N2A-N2B receptors were purified with Strep resin and analyzed by western blot. Representative blots for antibodies proved GluN2A (Strep) and GluN2B (Flag) from cells with different treatments as indicated on the top of this panel.

**Table s1.**
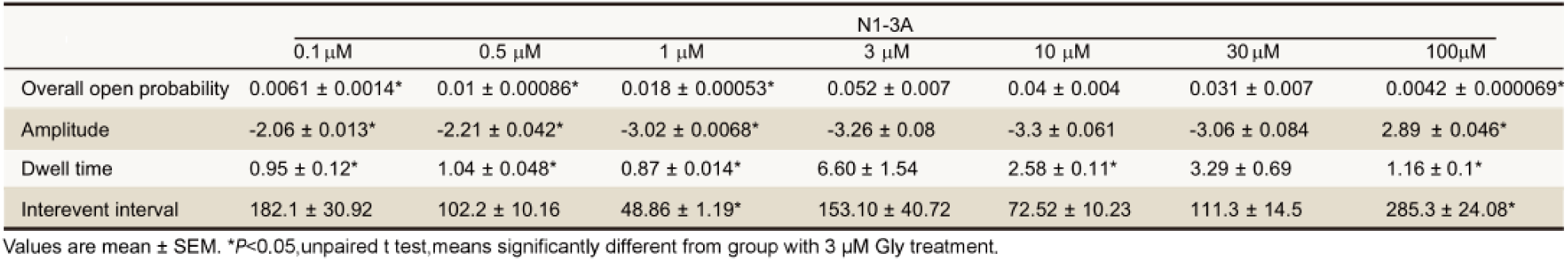
Single-Channel Properties of N1-N3A receptors.

Summary of Cryo-EM data collection and data analysis
| GluN1-2A-3A receptors<br>(Fabs/ Gly/Glu) |  |
| --- | --- |
| **Data collection and Processing** |  |
| Magnification |  |
| Voltage (kV) | 300 |
| Electron exposure (e-/Å<sup>2</sup>) | 60 |
| Defocus range (μm) | -1.5-3.0 |
| Pixel size (Å) | 1.08 |
| Symmetry imposed | C1 |
| Initial particle images (no.) | 528,709 |
| Final particle images (no.) | 162,395 |
| Map resolution (Å) | 7.73 |
| FSC threshold | 0.14 |
| **Refinement** |  |
| Initial model used (PDB code) | 6MMP |
| Model resolution (Å) | 6.88 |
| FSC threshold | 0.14 |
| **Model composition** |  |
| Non-hydrogen atoms | 30178 |
| Protein residues | 3974 |
| **Validation** |  |
| Mol Probity score | 1.98 |
| Clashscore | 7.49 |
| **Ramachandran plot** |  |
| Favored (%) | 96.64 |
| Allowed (%) | 3.36 |
| Disallowed (%) | 2.96 |

## Reference

1 Jalali-Yazdi, F., Chowdhury, S., Yoshioka, C. & Gouaux, E. Mechanisms for Zinc and Proton Inhibition of the GluN1/GluN2A NMDA Receptor. Cell 175, 1520–1532 e1515, doi:10.1016/j.cell.2018.10.043 (2018).

2 Lu, W., Du, J., Goehring, A. & Gouaux, E. Cryo-EM structures of the triheteromeric NMDA receptor and its allosteric modulation. Science 355, doi:10.1126/science.aal3729 (2017).

3 Zhang, Y. et al. Structural basis of ketamine action on human NMDA receptors. Nature 596, 301–305, doi:10.1038/s41586-021-03769-9 (2021).

4 Wang, H. et al. Gating mechanism and a modulatory niche of human GluN1-GluN2A NMDA receptors. Neuron 109, 2443–2456 e2445, doi:10.1016/j.neuron.2021.05.031 (2021).

5 Zhang, M. et al. Assembly and architecture of endogenous NMDA receptors in adult cerebral cortex and hippocampus. Cell, doi:10.1016/j.cell.2025.01.004 (2025).

6 Wang, H. et al. Structural basis for antibody-mediated NMDA receptor clustering and endocytosis in autoimmune encephalitis. Nature structural & molecular biology 31, 1987–1996, doi:10.1038/s41594-024-01387-3 (2024).

7 Zhang, J. et al. Distinct structure and gating mechanism in diverse NMDA receptors with GluN2C and GluN2D subunits. Nature structural & molecular biology 30, 629–639, doi:10.1038/s41594-023-00959-z (2023).

8 Liu, Y., Jia, Y. & Kou, Z. Bibliometric analysis of NMDA receptors: 2015–2024. Frontiers in pharmacology 16, doi:10.3389/fphar.2025.1614831 (2025).

9 Hurley, E. P. et al. GluN3A and Excitatory Glycine Receptors in the Adult Hippocampus. The Journal of neuroscience : the official journal of the Society for Neuroscience 44, doi:10.1523/JNEUROSCI.0401-24.2024 (2024).

10 Berg, L. K., Larsson, M., Morland, C. & Gundersen, V. Pre- and postsynaptic localization of NMDA receptor subunits at hippocampal mossy fibre synapses. Neuroscience 230, 139–150, doi:10.1016/j.neuroscience.2012.10.061 (2013).

11 Beesley, S., Gunjan, A. & Kumar, S. S. Visualizing the triheteromeric N-methyl-D-aspartate receptor subunit composition. Frontiers in synaptic neuroscience 15, 1156777, doi:10.3389/fnsyn.2023.1156777 (2023).

12 Wu, E., Zhang, J., Zhang, J. & Zhu, S. Structural insights into gating mechanism and allosteric regulation of NMDA receptors. Current opinion in neurobiology 83, 102806, doi:10.1016/j.conb.2023.102806 (2023).

13 Michalski, K. & Furukawa, H. Structure and function of GluN1-3A NMDA receptor excitatory glycine receptor channel. Sci Adv 10, eadl5952, doi:10.1126/sciadv.adl5952 (2024).

14 Xiong, K., Lou, S., Lian, Z., Wu, Y. & Kou, Z. The GluN3-containing NMDA receptors. Channels 19, 2490308, doi:10.1080/19336950.2025.2490308 (2025).

15 Liu, Y., Shao, D., Lou, S. & Kou, Z. Structural prediction of GluN3 NMDA receptors. Frontiers in physiology 15, 1446459, doi:10.3389/fphys.2024.1446459 (2024).

16 Perez-Otano, I., Larsen, R. S. & Wesseling, J. F. Emerging roles of GluN3-containing NMDA receptors in the CNS. Nature reviews. Neuroscience 17, 623–635, doi:10.1038/nrn.2016.92 (2016).

17 Yao, Y., Harrison, C. B., Freddolino, P. L., Schulten, K. & Mayer, M. L. Molecular mechanism of ligand recognition by NR3 subtype glutamate receptors. The EMBO journal 27, 2158–2170, doi:10.1038/emboj.2008.140 (2008).

18 Wong, H. K. et al. Temporal and regional expression of NMDA receptor subunit NR3A in the mammalian brain. The Journal of comparative neurology 450, 303–317, doi:10.1002/cne.10314 (2002).

19 Roberts, A. C. et al. Downregulation of NR3A-containing NMDARs is required for synapse maturation and memory consolidation. Neuron 63, 342–356, doi:10.1016/j.neuron.2009.06.016 (2009).

20 Kehoe, L. A. et al. GluN3A promotes dendritic spine pruning and destabilization during postnatal development. The Journal of neuroscience : the official journal of the Society for Neuroscience 34, 9213–9221, doi:10.1523/jneurosci.5183-13.2014 (2014).

21 Murillo, A. et al. Temporal Dynamics and Neuronal Specificity of Grin3a Expression in the Mouse Forebrain. Cerebral cortex 31, 1914–1926, doi:10.1093/cercor/bhaa330 (2021).

22 Larsen, R. S. et al. Synapse-specific control of experience-dependent plasticity by presynaptic NMDA receptors. Neuron 83, 879–893, doi:10.1016/j.neuron.2014.07.039 (2014).

23 Bossi, S. et al. GluN3A excitatory glycine receptors control adult cortical and amygdalar circuits. Neuron, doi:10.1016/j.neuron.2022.05.016 (2022).

24 Otsu, Y. et al. Control of aversion by glycine-gated GluN1/GluN3A NMDA receptors in the adult medial habenula. Science 366, 250–254, doi:10.1126/science.aax1522 (2019).

25 Das, S. et al. Increased NMDA current and spine density in mice lacking the NMDA receptor subunit NR3A. Nature 393, 377–381, doi:10.1038/30748 (1998).

26 Gonzalez-Gonzalez, I. M. et al. GluN3A subunit tunes NMDA receptor synaptic trafficking and content during postnatal brain development. Cell reports 42, 112477, doi:10.1016/j.celrep.2023.112477 (2023).

27 Larsen, R. S. et al. NR3A-containing NMDARs promote neurotransmitter release and spike timing- dependent plasticity. Nature neuroscience 14, 338–344, doi:10.1038/nn.2750 (2011).

28 Chen, L. F. et al. The NMDA receptor subunit GluN3A regulates synaptic activity-induced and myocyte enhancer factor 2C (MEF2C)-dependent transcription. The Journal of biological chemistry 295, 8613-8627, doi:10.1074/jbc.RA119.010266 (2020).

29 Jin, Z. et al. Selective increases of AMPA, NMDA, and kainate receptor subunit mRNAs in the hippocampus and orbitofrontal cortex but not in prefrontal cortex of human alcoholics. Frontiers in cellular neuroscience 8, 11, doi:10.3389/fncel.2014.00011 (2014).

30 Xie, X. et al. Association between genetic variations of NMDA receptor NR3 subfamily genes and heroin addiction in male Han Chinese. Neuroscience letters 631, 122–125, doi:10.1016/j.neulet.2016.08.025 (2016).

31 Salter, M. G. & Fern, R. NMDA receptors are expressed in developing oligodendrocyte processes and mediate injury. Nature 438, 1167–1171, doi:10.1038/nature04301 (2005).

32 Karadottir, R., Cavelier, P., Bergersen, L. H. & Attwell, D. NMDA receptors are expressed in oligodendrocytes and activated in ischaemia. Nature 438, 1162–1166, doi:10.1038/nature04302 (2005).

33 Micu, I. et al. NMDA receptors mediate calcium accumulation in myelin during chemical ischaemia. Nature 439, 988–992, doi:10.1038/nature04474 (2006).

34 Lee, J. H. et al. A neuroprotective role of the NMDA receptor subunit GluN3A (NR3A) in ischemic stroke of the adult mouse. American journal of physiology. Cell physiology 308, C570–577, doi:10.1152/ajpcell.00353.2014 (2015).

35 Marco, S. et al. Suppressing aberrant GluN3A expression rescues synaptic and behavioral impairments in Huntington’s disease models. Nature medicine 19, 1030–1038, doi:10.1038/nm.3246 (2013).

36 Mahfooz, K. et al. GluN3A promotes NMDA spiking by enhancing synaptic transmission in Huntington’s disease models. Neurobiology of disease 93, 47–56, doi:10.1016/j.nbd.2016.04.001 (2016).

37 Marco, S., Murillo, A. & Perez-Otano, I. RNAi-Based GluN3A Silencing Prevents and Reverses Disease Phenotypes Induced by Mutant huntingtin. Molecular therapy : the journal of the American Society of Gene Therapy 26, 1965–1972, doi:10.1016/j.ymthe.2018.05.013 (2018).

38 Zhong, W. et al. Pathogenesis of sporadic Alzheimer’s disease by deficiency of NMDA receptor subunit GluN3A. Alzheimer’s & dementia : the journal of the Alzheimer’s Association 18, 222–239, doi:10.1002/alz.12398 (2022).

39 Yuan, T. et al. Expression of cocaine-evoked synaptic plasticity by GluN3A-containing NMDA receptors. Neuron 80, 1025–1038, doi:10.1016/j.neuron.2013.07.050 (2013).

40 Pfisterer, U. et al. Identification of epilepsy-associated neuronal subtypes and gene expression underlying epileptogenesis. Nature communications 11, 5038, doi:10.1038/s41467-020-18752-7 (2020).

41 Kou, Z. Comment on the use of AI-based protein design for autoimmune encephalitis: exciting possibilities and practical considerations. Multiple sclerosis and related disorders 101, 106593, doi:10.1016/j.msard.2025.106593 (2025).

42 Zhang, M. et al. Dysfunction of GluN3A subunit is involved in depression-like behaviors through synaptic deficits. J Affect Disord 332, 72–82, doi:10.1016/j.jad.2023.03.076 (2023).

43 Chatterton, J. E. et al. Excitatory glycine receptors containing the NR3 family of NMDA receptor subunits. Nature 415, 793–798, doi:10.1038/nature715 (2002).

44 Madry, C. et al. Principal role of NR3 subunits in NR1/NR3 excitatory glycine receptor function. Biochemical and biophysical research communications 354, 102–108, doi:10.1016/j.bbrc.2006.12.153 (2007).

45 Nilsson, A. et al. Analysis of NR3A receptor subunits in human native NMDA receptors. Brain research 1186, 102–112, doi:10.1016/j.brainres.2007.09.008 (2007).

46 Bossi, S., Pizzamiglio, L. & Paoletti, P. Excitatory GluN1/GluN3A glycine receptors (eGlyRs) in brain signaling. Trends in neurosciences, doi:10.1016/j.tins.2023.05.002 (2023).

47 Auberson, Y. P. et al. N-phosphonoalkyl-5-aminomethylquinoxaline-2,3-diones: in vivo active AMPA and NMDA(glycine) antagonists. Bioorg Med Chem Lett 9, 249–254, doi:10.1016/s0960-894x(98)00720-3 (1999).

48 Rouzbeh, N. et al. Allosteric modulation of GluN1/GluN3 NMDA receptors by GluN1-selective competitive antagonists. The Journal of general physiology 155, doi:10.1085/jgp.202313340 (2023).

49 Madry, C., Betz, H., Geiger, J. R. & Laube, B. Supralinear potentiation of NR1/NR3A excitatory glycine receptors by Zn2+ and NR1 antagonist. Proceedings of the National Academy of Sciences of the United States of America 105, 12563–12568, doi:10.1073/pnas.0805624105 (2008).

50 Awobuluyi, M. et al. Subunit-specific roles of glycine-binding domains in activation of NR1/NR3 N- methyl-D-aspartate receptors. Molecular pharmacology 71, 112–122, doi:10.1124/mol.106.030700 (2007).

51 Grand, T., Abi Gerges, S., David, M., Diana, M. A. & Paoletti, P. Unmasking GluN1/GluN3A excitatory glycine NMDA receptors. Nature communications 9, 4769, doi:10.1038/s41467-018-07236-4 (2018).

52 Kang, H. et al. Structural basis for channel gating and blockade in tri-heteromeric GluN1-2B-2D NMDA receptor. Neuron, doi:10.1016/j.neuron.2025.01.013 (2025).

53 Perez-Otano, I. et al. Assembly with the NR1 subunit is required for surface expression of NR3A- containing NMDA receptors. The Journal of neuroscience : the official journal of the Society for Neuroscience 21, 1228–1237 (2001).

54 Sasaki, Y. F. et al. Characterization and comparison of the NR3A subunit of the NMDA receptor in recombinant systems and primary cortical neurons. Journal of neurophysiology 87, 2052–2063, doi:10.1152/jn.00531.2001 (2002).

55 Burzomato, V., Frugier, G., Perez-Otano, I., Kittler, J. T. & Attwell, D. The receptor subunits generating NMDA receptor mediated currents in oligodendrocytes. The Journal of physiology 588, 3403–3414, doi:10.1113/jphysiol.2010.195503 (2010).

56 Tong, G. et al. Modulation of NMDA receptor properties and synaptic transmission by the NR3A subunit in mouse hippocampal and cerebrocortical neurons. Journal of neurophysiology 99, 122–132, doi:10.1152/jn.01044.2006 (2008).

57 Al-Hallaq, R. A. et al. Association of NR3A with the N-methyl-D-aspartate receptor NR1 and NR2 subunits. Molecular pharmacology 62, 1119–1127 (2002).

58 Chou, T. H., Tajima, N., Romero-Hernandez, A. & Furukawa, H. Structural Basis of Functional Transitions in Mammalian NMDA Receptors. Cell 182, 357–371 e313, doi:10.1016/j.cell.2020.05.052 (2020).

59 Yao, Y., Belcher, J., Berger, A. J., Mayer, M. L. & Lau, A. Y. Conformational analysis of NMDA receptor GluN1, GluN2, and GluN3 ligand-binding domains reveals subtype-specific characteristics. Structure (London, England : 1993) 21, 1788-1799, doi:10.1016/j.str.2013.07.011 (2013).

60 Yao, Z. et al. A taxonomy of transcriptomic cell types across the isocortex and hippocampal formation. Cell 184, 3222–3241 e3226, doi:10.1016/j.cell.2021.04.021 (2021).

61 Guo, C., Rudolph, S., Neuwirth, M. E. & Regehr, W. G. Purkinje cell outputs selectively inhibit a subset of unipolar brush cells in the input layer of the cerebellar cortex. eLife 10, doi:10.7554/eLife.68802 (2021).

62 Mickelsen, L. E. et al. Cellular taxonomy and spatial organization of the murine ventral posterior hypothalamus. eLife 9, doi:10.7554/eLife.58901 (2020).

63 O’Leary, T. P. et al. Extensive and spatially variable within-cell-type heterogeneity across the basolateral amygdala. eLife 9, doi:10.7554/eLife.59003 (2020).

64 Zhang, J. et al. Distinct structure and gating mechanism in diverse NMDA receptors with GluN2C and GluN2D subunits. Nature structural & molecular biology, doi:10.1038/s41594-023-00959-z (2023).

65 Wyllie, D. J., Behe, P. & Colquhoun, D. Single-channel activations and concentration jumps: comparison of recombinant NR1a/NR2A and NR1a/NR2D NMDA receptors. The Journal of physiology 510 **(Pt** **1**), 1–18, doi:10.1111/j.1469-7793.1998.001bz.x (1998).

66 Zhu, S. et al. Genetically encoding a light switch in an ionotropic glutamate receptor reveals subunit- specific interfaces. Proceedings of the National Academy of Sciences of the United States of America 111, 6081–6086, doi:10.1073/pnas.1318808111 (2014).

67 Ye, S., Huber, T., Vogel, R. & Sakmar, T. P. FTIR analysis of GPCR activation using azido probes. Nature chemical biology 5, 397–399, doi:10.1038/nchembio.167 (2009).

68 ., Wang, L., Brock, A., Herberich, B. & Schultz, P. G. Expanding the genetic code of Escherichia coli. Science 292, 498–500, doi:10.1126/science.1060077 (2001).

69 Poulsen, M. H., Poshtiban, A., Klippenstein, V., Ghisi, V. & Plested, A. J. R. Gating modules of the AMPA receptor pore domain revealed by unnatural amino acid mutagenesis. Proceedings of the National Academy of Sciences of the United States of America, doi:10.1073/pnas.1818845116 (2019).

70 Koole, C. et al. Genetically encoded photocross-linkers determine the biological binding site of exendin-4 peptide in the N-terminal domain of the intact human glucagon-like peptide-1 receptor (GLP-1R). The Journal of biological chemistry 292, 7131-7144, doi:10.1074/jbc.M117.779496 (2017).

71 Kang, J. Y. et al. In vivo expression of a light-activatable potassium channel using unnatural amino acids. Neuron 80, 358–370, doi:10.1016/j.neuron.2013.08.016 (2013).

72 Luo, J., Wang, Y., Yasuda, R. P., Dunah, A. W. & Wolfe, B. B. The majority of N-methyl-D-aspartate receptor complexes in adult rat cerebral cortex contain at least three different subunits (NR1/NR2A/NR2B). Molecular pharmacology 51, 79–86, doi:10.1124/mol.51.1.79 (1997).

73 Liu, Y., Tang, H., Zhang, J., Li, D. & Kou, Z. Artificial intelligence insight on structural basis and small molecule binding niches of NMDA receptor. Comput Struct Biotechnol J **27**, 3167-3180, doi:10.1016/j.csbj.2025.07.027 (2025).

74 Hansen, K. B., Ogden, K. K., Yuan, H. & Traynelis, S. F. Distinct functional and pharmacological properties of Triheteromeric GluN1/GluN2A/GluN2B NMDA receptors. Neuron 81, 1084–1096, doi:10.1016/j.neuron.2014.01.035 (2014).

75 Bhattacharya, S. et al. Triheteromeric GluN1/GluN2A/GluN2C NMDARs with Unique Single-Channel Properties Are the Dominant Receptor Population in Cerebellar Granule Cells. Neuron 99, 315–328 e315, doi:10.1016/j.neuron.2018.06.010 (2018).

76 Yi, F., Bhattacharya, S., Thompson, C. M., Traynelis, S. F. & Hansen, K. B. Functional and pharmacological properties of triheteromeric GluN1/2B/2D NMDA receptors. The Journal of physiology, doi:10.1113/JP278168 (2019).

77 Rauner, C. & Kohr, G. Triheteromeric NR1/NR2A/NR2B receptors constitute the major N-methyl-D- aspartate receptor population in adult hippocampal synapses. The Journal of biological chemistry 286, 7558–7566, doi:10.1074/jbc.M110.182600 (2011).

78 Stroebel, D., Carvalho, S., Grand, T., Zhu, S. & Paoletti, P. Controlling NMDA receptor subunit composition using ectopic retention signals. The Journal of neuroscience : the official journal of the Society for Neuroscience 34, 16630–16636, doi:10.1523/JNEUROSCI.2736-14.2014 (2014).

79 Kang, H. et al. Structural basis for channel gating and blockade in tri-heteromeric GluN1-2B-2D NMDA receptor. Neuron 113, 991–1005 e1005, doi:10.1016/j.neuron.2025.01.013 (2025).

80 Ciabarra, A. M. et al. Cloning and characterization of chi-1: a developmentally regulated member of a novel class of the ionotropic glutamate receptor family. The Journal of neuroscience : the official journal of the Society for Neuroscience 15, 6498–6508 (1995).

81 Jia, Y. et al. TMC1 and TMC2 Proteins Are Pore-Forming Subunits of Mechanosensitive Ion Channels. Neuron 105, 310–321 e313, doi:10.1016/j.neuron.2019.10.017 (2020).

82 Bessa-Neto, D. et al. Bioorthogonal labeling of transmembrane proteins with non-canonical amino acids unveils masked epitopes in live neurons. Nature communications 12, 6715, doi:10.1038/s41467-021-27025-w (2021).

83 Arsic, A., Hagemann, C., Stajkovic, N., Schubert, T. & Nikic-Spiegel, I. Minimal genetically encoded tags for fluorescent protein labeling in living neurons. Nature communications 13, 314, doi:10.1038/s41467-022-27956-y (2022).

84 Mulhall, E. M. et al. Direct observation of the conformational states of PIEZO1. Nature, doi:10.1038/s41586-023-06427-4 (2023).

85 Zhao, Y., Chen, S., Swensen, A. C., Qian, W. J. & Gouaux, E. Architecture and subunit arrangement of native AMPA receptors elucidated by cryo-EM. Science, doi:10.1126/science.aaw8250 (2019).

86 Lee, C. H. et al. NMDA receptor structures reveal subunit arrangement and pore architecture. Nature 511, 191–197, doi:10.1038/nature13548 (2014).

87 Scheres, S. H. RELION: implementation of a Bayesian approach to cryo-EM structure determination. J Struct Biol 180, 519–530, doi:10.1016/j.jsb.2012.09.006 (2012).

88 Zheng, S. Q. et al. MotionCor2: anisotropic correction of beam-induced motion for improved cryo- electron microscopy. Nat Methods 14, 331–332, doi:10.1038/nmeth.4193 (2017).

89 Zhang, K. Gctf: Real-time CTF determination and correction. J Struct Biol 193, 1–12, doi:10.1016/j.jsb.2015.11.003 (2016).

90 Arnold, K., Bordoli, L., Kopp, J. & Schwede, T. The SWISS-MODEL workspace: a web-based environment for protein structure homology modelling. Bioinformatics 22, 195–201, doi:10.1093/bioinformatics/bti770 (2006).

91 Pettersen, E. F. et al. UCSF ChimeraX: Structure visualization for researchers, educators, and developers. Protein Sci 30, 70–82, doi:10.1002/pro.3943 (2021).

92 Emsley, P. & Cowtan, K. Coot: model-building tools for molecular graphics. Acta Crystallogr D Biol Crystallogr 60, 2126–2132, doi:10.1107/S0907444904019158 (2004).

93 Afonine, P. V. et al. Real-space refinement in PHENIX for cryo-EM and crystallography. Acta Crystallogr D Struct Biol 74, 531–544, doi:10.1107/S2059798318006551 (2018).

94 Kucukelbir, A., Sigworth, F. J. & Tagare, H. D. Quantifying the local resolution of cryo-EM density maps. Nat Methods 11, 63–65, doi:10.1038/nmeth.2727 (2014).

95 Wu, K. W., Kou, Z. W., Mo, J. L., Deng, X. X. & Sun, F. Y. Neurovascular coupling protects neurons against hypoxic injury via inhibition of potassium currents by generation of nitric oxide in direct neuron and endothelium cocultures. Neuroscience 334, 275–282, doi:10.1016/j.neuroscience.2016.08.012 (2016).

96 Kou, Z. W. et al. Vascular endothelial growth factor increases the function of calcium-impermeable AMPA receptor GluA2 subunit in astrocytes via activation of protein kinase C signaling pathway. Glia 67, 1344–1358, doi:10.1002/glia.23609 (2019).

97 Meyer-Dilhet, G. & Courchet, J. In Utero Cortical Electroporation of Plasmids in the Mouse Embryo. STAR Protoc 1, 100027, doi:10.1016/j.xpro.2020.100027 (2020).

98 Mo, J. L. et al. MicroRNA-365 modulates astrocyte conversion into neuron in adult rat brain after stroke by targeting Pax6. Glia 66, 1346–1362, doi:10.1002/glia.23308 (2018).

99 Selvakumar, P. et al. Structural and compositional diversity in the kainate receptor family. Cell reports 37, 109891, doi:10.1016/j.celrep.2021.109891 (2021).

